# Computational investigation reveals that the mutant strains of SARS-CoV2 are highly infectious than wildtype

**DOI:** 10.1101/2021.04.23.441125

**Authors:** Rakesh Kumar, Rahul Kumar, Harsh Goel, Pranay Tanwar

**Affiliations:** Dr.B.R.A.-Institute Rotary Cancer Hospital, All India Institute of Medical Sciences, New Delhi, INDIA-110029

**Keywords:** SARS-CoV2, mutant strains, molecular dynamics, network analysis, protein-protein interaction

## Abstract

Remarkable infectivity of severe acute respiratory syndrome-coronavirus 2 (SARS-CoV2) is due to the rapid emergence of various strains, thus enable the virus to rule the world. Over the course of SARS-CoV2 pandemic, the scientific communities worldwide are responding to newly emerging genetic variants. However, the mechanism behind the persistent infection of these variants is still not known due to the paucity of study of these variants at molecular level. In this scenario, computational methods have immense utility in understanding the molecular and functional properties of different variants. Therefore, in this study various mutants (MTs) of SpikeS1 receptor binding domain (RBD) of highly infectious SARS-CoV2 strains were carried and elucidated the protein structure and dynamics using molecular dynamics (MD) approach. MD simulation study showed that all MTs exhibited stable structures with altered functional properties. Furthermore, the binding strength of different MTs along with WT (wildtype) was revealed through protein-protein docking and observed that MTs showed high binding affinities than WT. Hence, this study shed light on the molecular basis of infection caused by different variants of SARS-CoV2, which might play an important role in to cease the transmission and pathogenesis of virus and also implicate in rational designing of a specific drug.

## 1. INTRODUCTION

Near the end of 2019, new public health crises arise from Wuhan city of China due to the outbreak of nCoV (novel coronavirus) or SARS-CoV2, which spread worldwide and named COVID-19 (coronavirus disease 2019) (Wang et al., 2020). Till now (February 2021), the cumulative number of total confirmed cases and deaths of COVID-19 reached up to ∼115 and ∼2 million (https://www.worldometers.info/coronavirus/), respectively (Fig. S1A&B). During the first wave of SARS-CoV2, most of the confirmed cases are observed in the USA (∼2.64 millions) (Fig. S1C; Table S1), while during the second wave, its number reached ∼20.06 million (Fig. S1E) (Roser et al., 2020). To date, ∼28.41 million confirmed cases were observed in the USA alone, which are the highest number of cases arisen in any country in entire world (Roser et al., 2020). However, the cumulative confirmed COVID-19 deaths during first (∼575.02 per million deaths) and second waves (∼1,226.54 deaths per million) of infections were reported highest in Italy (Fig. S1D&F) (Bontempi, 2021). While in UK, ∼1,801.49 deaths per million were observed, which are the highest number of deaths per million estimated by any country throughout the world (Table S2). Rapid global spread and transmission of COVID-19 provide the virus with substantial opportunities for the natural selection of favourable mutations (Wu et al., 2020). Mutation variants in SARS-CoV2 genomes have raised serious concerns about changes in infectivity (Young et al., 2020). The emergence of new strains driven by the adaptive mutation indicates its instability over a period of time and can contribute towards its virulence. This rapid evolution seems to be a common problem that shows the gravity of coronavirus infection as a communicable disease and attract the attention of the scientific community as a matter of serious concern worldwide (Menachery et al., 2016). Mostly, different strains are arising due to mutations in the receptor binding domain (RBD) of SpikeS1 protein (Korber et al., 2020). SpikesS1 protein of virus interacts with host angiotensin converting enzyme2 (ACE2), thus enable the virus to enter into the host cell (Yang et al., 2020). V367F (Valine 367 to Phenylalanine) mutations in the RBD of SpikeS1 protein was reported from the strains found in Wuhan, Shenzhen, Hong Kong and France (Starr et al., 2020; Tang et al., 2020). R408I (Arginine 408 to Isoleucine) mutant isolates from India (Saha et al., 2020). In addition, mutations such as G476S (Glycine 476 to Serine) and V483A (Valine 483 to Alanine) are reported in the United States (Wang et al., 2021). Recently, a new strain N501Y in which Asparagine positioned at 501 of RBD is substituted by Tyrosine arises in United Kingdom and South Africa (Tegally et al., 2020; Wise, 2020). These mutations (V367F, R408I, G476S, V483A and N501Y) were mainly located in the SpikeS1 RBD domain. Of these five, three mutations like G476S, V483A and N501Y occur at the binding interface of RBD (C-terminal side) and PD (peptidase domain) of the host ACE2 receptor (Lan et al., 2020). The rest of mutations V367F and R408I, mostly affected the overall topology and stability of RBD as these mutations present at the loop regions of N-terminal and turn that connect β-sheet3 and 4, respectively (Yan et al., 2020). These mutations (V367F, R408I, G476S, V483A and N501Y) indirectly assist in the stable binding of RBD to the host ACE2 receptor (Lan et al., 2020). In addition to the above mutations, there are various other mutations found in the SpikeS1 protein beyond the RBD domain that also contributed to the virus infectivity (Yan et al., 2020). Since targeting the mutations in RBD domain provide the first step to halt the virus infection, therefore the current study is mainly focused on the RBD mutations. To unravel the stability and dynamics of highly infectious strains or RBD mutants (MTs), long run molecular dynamics (MD) simulations were performed followed by elucidating the key residues involved in signalling through residues interaction network (RIN) approach. Furthermore, the infectiveness of above MTs were assessed through protein-protein (ACE2-RBD) docking of both WT and MT RBD proteins. We observed that all MTs showed similar dynamic but differential structural properties. Moreover, RBD MTs showed maximum binding affinities to the host ACE2 receptor protein. Hence, the work in the present study will help in the creation of a more specific inhibitor or drug by taking the above residues in consideration.

## 2. MATERIALS AND METHODS

### 2.1 3D structures preparation and validation

The tertiary structure of RBD of Spike S1 protein was taken from RCSB-PDB (Research collaboration for structural bioinformatics-protein data bank), which was experimentally resolved in complexed form (PDB id: 6M17) (Rose et al., 2011; Yan et al., 2020). The structure was prepared in PyMOL (The PyMOL Molecular Graphics System, Version 1.3 Schrodinger, LLC) and all other structures except RBD were excluded. Distinct missense mutations which represented different strains of SARS-CoV2 such as V367F (Valine to Phenylalanine), R408I (Arginine to Isoleucine), G476S (Glycine to Serine), V483A (Valine to Alanine), N501Y (Asparagine to Tyrosine) were prepared in PyMOL as previously described (Ningombam et al., 2021). 3D structures of all mutants (V367F, R408I, G476S, V483A and N501Y) along with wildtype were energy minimised by using GROMACS (Van Der Spoel et al., 2005). Energy minimised protein models were subjected to validation by different quality estimation tools. Stereochemical properties, as well as local and global qualities of all models were monitored through PROCHECK module of SAVES (structure analysis and verification server), ProSA (Protein structure analysis; https://prosa.services.came.sbg.ac.at/prosa.php) and QMEAN (Qualitative model energy analysis; https://swissmodel.expasy.org/qmean/) web servers, respectively (Benkert et al., 2008; Luthy et al., 1992; Wiederstein & Sippl, 2007).

### 2.2 Molecular dynamics simulation

Molecular dynamics (MD) simulations of 150ns for each WT and MTs were conducted using GROMACS in conjunction with Amber force field (Oostenbrink et al., 2004; Van Der Spoel et al., 2005). Initially, GROMACS readable coordinates and topology files were generated through “pdb2gmx” module and all systems (WT and MTs) were solvated using “editconf” and solvate scripts. Systems solvation were accomplished by immersing all WT and MT proteins into triclinic box with TIP3P water model. After that, all systems were neutralized by adding sodium and chloride counterions using “genion” module followed by energy minimisation through the steepest-descent method. Hence after, two equilibration steps as NVT (constant number of particles, volume, and temperature) and NPT (constant number of particles, pressure, and temperature) ensembles were performed for 100 and 500ps at a temperature of 300K and pressure of 1bar, respectively. The temperature and pressure of all the systems were carried by v-rescale (a modified Berendsen method) and Parrinello-Rahman methods, respectively. Long range electrostatic interactions were preserved by using Particle Mesh Ewald (PME) methods and all bonds were constraint using Linear constraint solver (LINCS) algorithm. Finally, the production run of 150ns was performed in which time steps of 2fs were applied. Resultant MD trajectories were analysed by using gmx energy, gmx rms, gmx rmsf, gmx gyrate, gmx sasa and gmx hbond modules of GROMACS. Secondary structures analysis was done through the dictionary of secondary structure of protein (DSSP) approach using do_dssp script (Kabsch & Sander, 1983).

## 2.3 Essential dynamics

Essential dynamics (ED) is an important parameter, used to generate coordinates of subspace in which the motion of a protein is likely to take place that determines its biological function (Amadei et al., 1993). A series of eigenvectors with respective eigenvalues were generated after the construction and diagonalization of the covariance matrix of backbone atoms from all WT and MT systems. We restricted our interpretation with backbone atoms in order to avoid statistical noise and utilised stabled trajectories of last 25ns of all WT and MT systems for our ED analysis. Simultaneously, to avoid the biologically misinterpretation of PCs we calculated cosine values of first three PCs that should be ≤0.2 (Hess, 2000). First three principle components (PC1, PC2 and PC3) or eigenvectors were used which usually accompanied with maximum motions that are biologically significant (Kumar & Saran, 2018). ED analysis was performed by gmx covar, gmx analyze and gmx anaeig modules of GROMACS.

### 2.4 Residues network analysis

Residues interacting networks (RIN) of WT and MTs were predicted on MD optimised protein structure using NAPS (Network Analysis of Protein Structure) webserver (Chakrabarty, & Parekh, 2016). The server facilitated qualitative and quantitative analyses of RIN and protein contact network was constructed based on physiochemical properties of residues in which different centralities were measured. We restrict our analysis with Cα atoms where the residue is treated as node with an unweighted edge drawn within distance of 7Å connecting other residues in the given network. NAPS administered different centrality measures, such as degree (C_D_), closeness (C_C_), and betweenness (C_B_). C_D_ is defined as the number of node link with the central node as mentioned in equation (1), while C_C_ is the average farness from the central node to other nodes within the network as mentioned in equations (1) and (2). C_B_ is depending upon the shortest path between the nodes. It is the ratio of shortest routes from one node to other passes through *u* and the entire shortest path within the network as described in equation (3).

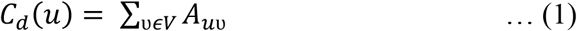

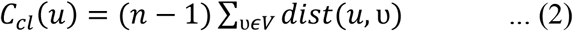

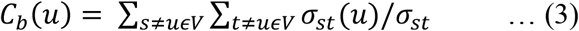

Whereas, A = adjacency matrix; V= set of all nodes; N= number of nodes in the network; dist (*u υ*) = shortest path distance between nodes u and *υ* σst = shortest path between s and t; σst (u) = shortest path between s and t passing through u.

### 2.5 Protein-protein docking

Protein-protein (p-p) docking of ACE2 and all WT and MTs SpikeS1 RBD were executed through HADDOCK (High Ambiguity Driven protein-protein DOCKing) webserver (van Zundert et al., 2016). Active residues which were identified through available experimental complex structures from PDB, used as constraint to generate AIRs (Ambiguous Interaction Restraints) during docking (Yan et al., 2020). HADDOCK carried p-p docking in 3 successive stages as previously described (Kumar & Mukherjee, 2021). Briefly, energy minimization steps were performed initially, followed by simulated annealing refinement and finally MD refinement with solvent. P-p complex structures obtained from the final stage were clustered and HADDOCK scores were calculated as given in equations (4-6).

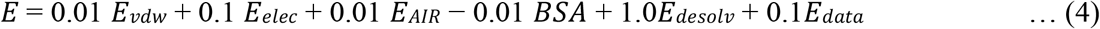

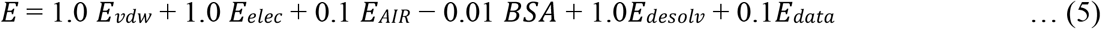

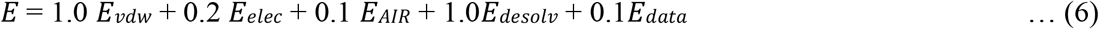

In the above equations, *E*_*vdw*_, *E*_*elec*_, *E*_*AIR*_, *E*_*desolv*_, *E*_*data*_ represented van der Waals, electrostatic, AIR, desolvation energies and energy of other restraint data and buried surface area, respectively. Best docked complexes with the lowest HADDOCK scores from all ACE-WT and -MT complexes were finally analysed. Protein-protein interaction visualisations at 3D and 2D levels and H-bonding were analysed in PyMOL (The PyMOL Molecular Graphics System, Version 1.3 Schrodinger, LLC) and DIMPLOT module of LigPlot+ v2.2.4, respectively (Laskowski & Swindells, 2011). Furthermore, the binding strength of various ACE2-WT and -MTs complexes were also verified by using mCSM-PPI2 server (Rodrigues et al., 2019). mCSM-PPI2 is novel computational tool that accurately predicts the impacts of missense mutations in p-p binding affinity. It used graph-based structural signatures (mCSM) to represent the wild-type residue environment, inter-residue non-covalent interaction networks, and evolutionary information, complex network metrics, and energetic terms to generate an optimized predictor.

## 3. RESULTS

### 3.1 WT and MTs demonstrated similar dynamic behaviours

The tertiary structure of RBD comprised about 181 amino acids long polypeptide and having seven β-sheets (Fig. 1A). An extended loop was present near C-terminal that facilitated its interaction with protein partner (ACE2). Various MT models were generated in PyMOL by replacing Valine367 with Phenylalanine (V367F), Arginine408 with Isoleucine (R408I), Glutamine476 with Serine (G476S), Valine483 with Alanine (V483A) and Asparagine501 with Tyrosine (N501Y) in WT 3D structure. WT and MT model structures were energy minimised and found that all models were exhibited the lowest values of minimum energies (data not shown). V367F and R408I MTs were present in between β-sheets 1 and 2, β-sheet 3 and 4, respectively (Fig. 1A). Additionally, an extended loop was present near to C-terminal harboured 2 mutations such as G476S and V483A, while N501Y mutation was located near to the end of C-terminal. The extended loop and the region near C-terminal of SpikeS1 RBD interacts with the peptidase domain (PD) of ACE receptor^15^. The structure validations of WT and newly constructed MT models were inspected by measuring the Phi/Psi angles in the Ramachandran plot and found that all 3D structures exhibited 68.5%, 29.5% and 2% of residues to be placed in favourable, allowed and disallowed regions, respectively (Table S3). Moreover, ProSA and QMEAN servers were also executed to analyse the local and global geometry of WT and MT structures and found that all WT and MT proteins exhibited ProSA and QMEAN values ranging from −5.15 to −5.25 and −7.33 to −8.15, respectively (Table S3). The negligible number of residues in disallowed regions and lowest values of ProSA and QMEAN scores indicated that all WT and MT protein structures had better stereochemical and geometrical properties.

**Fig. 1.**
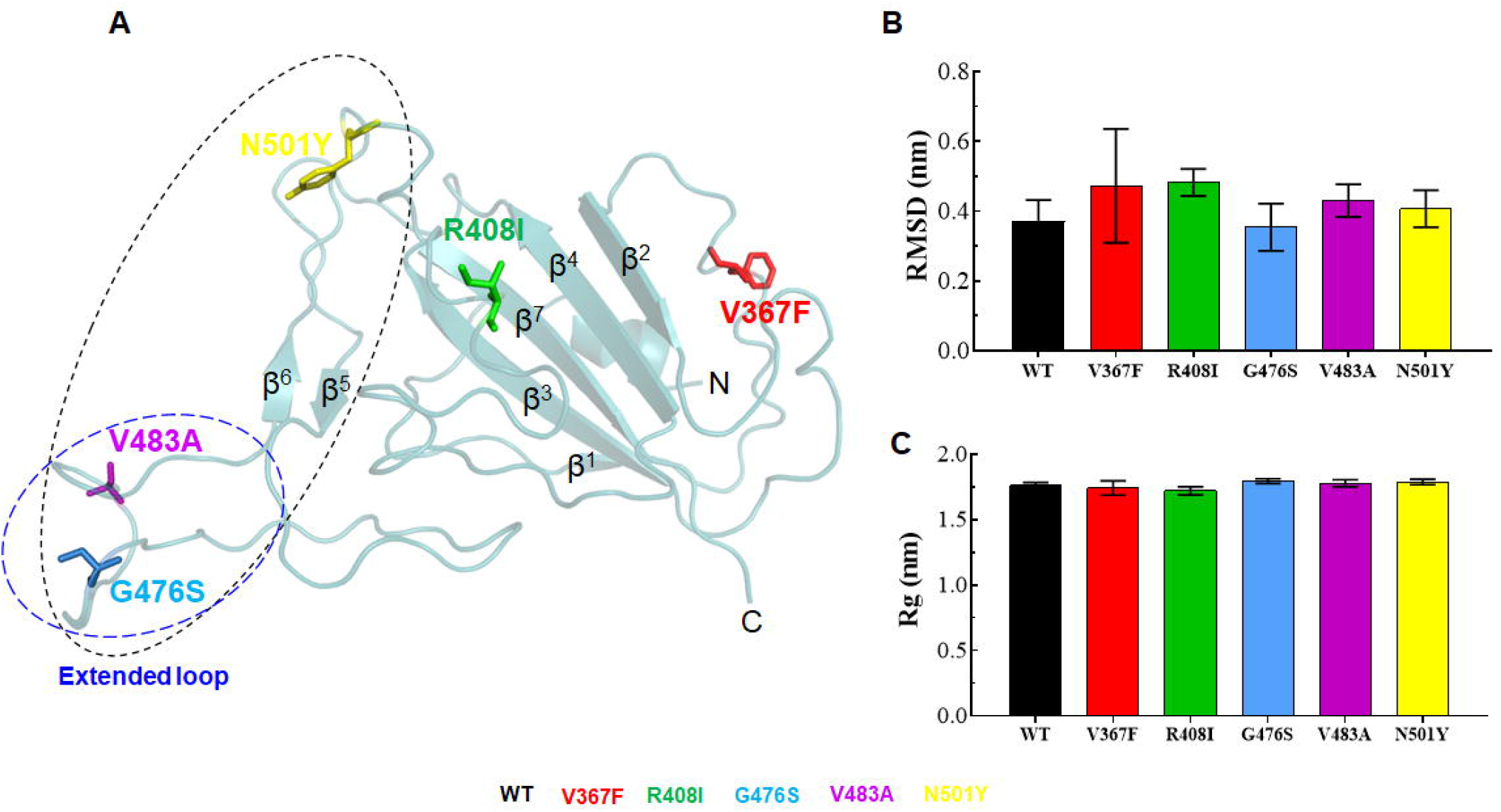
Tertiary structure, RMSD and Rg analyses. (A) Tertiary structure of SpikeS1 RBD of SARS-CoV2, (B) Root mean square deviation at function of time and (C) Radius of gyration at function of time in nanoseconds. 3D structure of RBD was shown in cartoon mode with cyan colour and different mutations V367F, R408I, G476S, V483A and N501Y were depicted in stick mode. Region interacting with host ACE2 of SpikeS1 RBD and extended loop were highlighted in back and blue dotted circles, respectively. Different sheets were labelled as β_1_-β_7_. WT, V367F, R408I, G476S, V483A and N501Y MTs were labelled in black, red, green, blue, magenta and yellow colour, respectively.

**Fig. 2.**
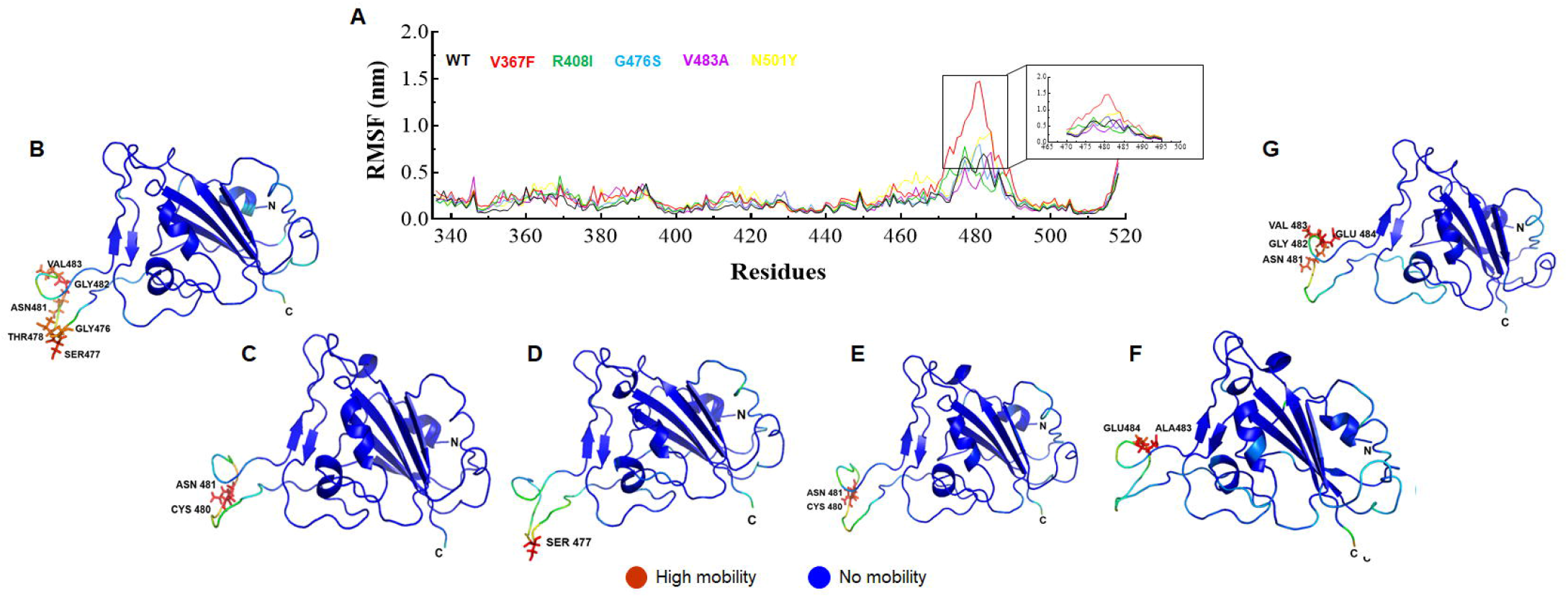
Root mean square fluctuation (RMSF) analysis. (A) RMSF plot at function of amino acid residues, (B) Structure fluctuations of WT, (C) Structure fluctuations of V367F MT, (D) Structure fluctuations of R408I MT, (E) Structure fluctuations of G476S MT, (F) Structure fluctuations of V483A MT and (G) Structure fluctuation of N501Y MT. In 2D RMSF plot, highly fluctuated regions were highlighted and static or mobile regions were shown as blue and red colour, respectively. Amino acid residues were depicted in stick mode with 3-letter code. WT, V367F, R408I, G476S, V483A and N501Y MTs were labelled in black, red, green, blue, magenta and yellow colour, respectively.

Preliminary analyses of effects of different mutations on 3D structure were predicted through MUpro (Cheng et al., 2006) and I-mutant (Capriotti et al., 2005) servers and found that all mutant except N501Y showed decreased structure stabilities (Tables S4 & S5). Further, detailed structural stability and dynamics were examined through MD simulations. To assess the structural stabilities of WT and MTs (V367F, R408I, G476S, V483A and N501Y), root mean square deviation (RMSD) of protein backbone with respect to equilibrium structures were calculated and found that average RMSD values of ∼0.37 for WT, ∼0.47 for V367F, ∼0.48 for R408I, ∼0.35 for G476S, ∼0.43 for V483A and ∼0.40nm for N501Y MTs were existed (Fig.1B). Further, RMSDs of all WT and MTs were equilibrated after 100ns time and showed similar pattern (Fig. S2A). Moreover, V367F and R408I MTs displayed higher deviations, whereas G476S MT exhibited lower deviation then WT, V483A and N501Y MTs (Fig.1B). RMSD of all MTs and WT showed a similar and consistent pattern. Protein compactness or globularity was measured by employing the radius of gyration (Rg) and found that average Rg values of ∼1.76, ∼1.74, ∼1.72, ∼1.79, ∼1.77 and ∼1.78nm were obtained for WT, V367F, R408I, G476S, V483A and N501Y MTs, respectively (Fig.1C). Rgs were slightly reduced in V367F and R408I MTs, whereas G476S, V483A and N501Y MTs showed slightly higher Rg then WT. Further, the Rg patterns of all MTs and WT were stabilised after ∼125ns time and showed similar behaviour (Fig. S2B). Taken together, all MTs and WT showed similar pattern of RMSD and Rg, indicating that both WT and MT proteins exhibited similar dynamic behaviours.

### 3.2 Root mean square fluctuations analyses of WT and MTs

Flexibility and stability of both WT and MTs were further accomplished by measuring root mean square fluctuation (RMSF) at residues level. RMSF is a crucial parameter use to monitor the structural flexibility due to side chain of residues. All MTs and WT showed mean RMSFs in the range of 0.14 to 0.2, excluding extended loop segment WT, V367F, R408I, G476S, V483A and N501Y MTs, respectively (Fig.2A). The extended loop segment present at the C-terminal side displayed higher fluctuation as compared to rest of the protein structures. It comprised ∼26 amino acid long segment (470-495) showed an average RMSFs of ∼0.37, ∼0.70, ∼0.39, ∼0.38, ∼0.32 and ∼0.46nm for WT, V367F, R408I, G476S, V483A and N501Y MTs, respectively (Fig.2A). V367F, R408I and N501Y MTs displayed slightly higher RMSFs, whereas V483A MT displayed lower RMSF then WT, indicating that V367F, R408I and N501Y MTs were more flexible and V483A MT was less flexible or more rigid than WT. Residues level inspection showing that Gly476, Ser477, Thr478, Asn481, Gly482 and Val483 remained highly mobile in WT (Fig.2B) whereas Cys480, Asn481 and Gly482 in V367F MT (Fig.2C), Ser477 in R408I (Fig.2D), Cys480 and Asn481 in G476S (Fig.2E), Ala483 and Glu484 in V483A and Asn481, Gly482 (Fig.2F), Val483 and Glu484 in N501Y MT (Fig.2G) residues remained to be highly mobile. RMSF results indicating that fluctuations were more pronounced at the loop region in all MTs and WT.

### 3.3 WT and MTs showed differential structural properties

Different structural properties of both WT and MT proteins were assessed through measuring the solvent accessible surface area (SASA), quantitative analysis of intra and inter hydrogen-bonding (H-bond) and qualitative as well as quantitative analyses of secondary structures formation during the entire simulation period. SASA was calculated through gmx sasa module of GROMACS and revealed that total SASA was higher in all MTs as compared to WT. An average SASA (total) values of WT, V367F, R408I, G476S, V483A and N501Y MTs were 103.2, 106.8, 106, 104.6, 108.7 and 108.3nm^2^, respectively (Fig.3A). Moreover, SASA behaviours remained constant and consistent during an entire simulation period in almost both WT and MTs (Fig. S3). During SASA analysis, hydrophobic remained slightly higher than the hydrophilic SASA in all WT and MT cases. Average values of hydrophobic SASA were lied in the range of 54.2nm^2^ for WT (low) to 58nm^2^ for V483A (high). While hydrophilic SASA values were in the range of 49nm^2^ for WT (low) to 51.5nm^2^ for N501Y (high) (Fig. 3A). This is quite possible because RBD of SpikeS1 formed a core of 7 β-Sheets which is largely hydrophobic in nature.

**Fig. 3.**
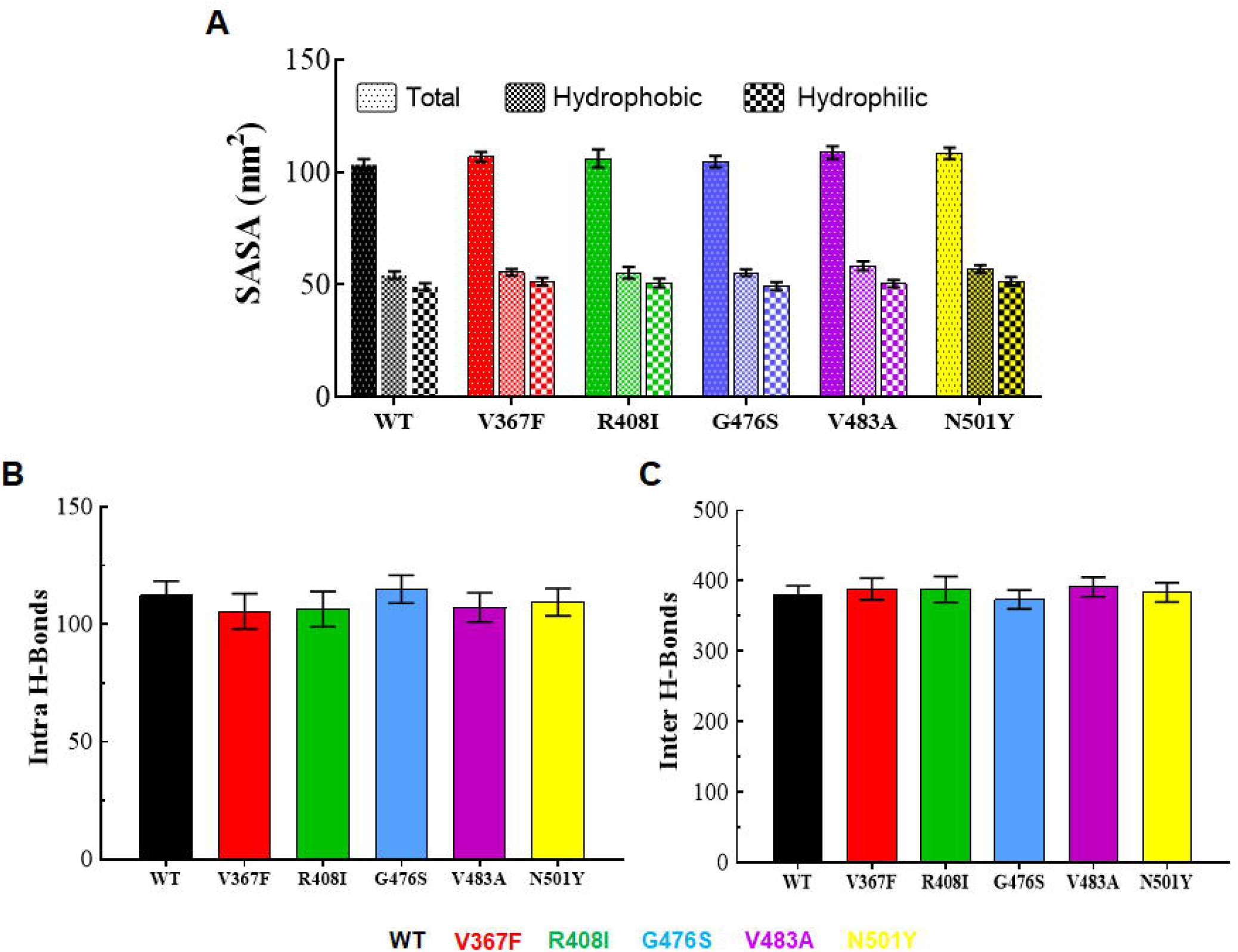
Structural properties analyses of WT and MT proteins. (A) Solvent accessible surface area (SASA) plot, (B) Protein-protein hydrogen bonds (H-bond) and (C) Protein-water hydrogen bonds (H-bond). WT, V367F, R408I, G476S, V483A and N501Y MTs were labelled in black, red, green, blue, magenta and yellow colour, respectively.

Structure integrity and interaction capacity of protein determined by various interaction forces out of which hydrogen bond (H-bond) plays a significant role in protein stability and functionality. We calculated both intra (protein-protein) and inter (protein-solvent) H-bond through gmx hbond module of GROMACS. WT, V367F, R408I, G476S, V483A, and N501Y MTs exhibited an average of 112.3, 105.5, 106.4, 115, 107.1, and 109.4 intra H-Bonds, respectively (Fig. 3B), while an average of 378.9, 388.5, 387.8, 373.4, 391.2, and 383.6 H-bonds of WT, V367F, R408I, G476S, V483A and N501Y were formed with the solvent molecule, respectively (Fig. 3C). Moreover, both intra and inter H-bonds remained consistent during an entire simulation period in WT and MT proteins (Fig. S4). During H-bond analysis, we observed that MTs exhibited high number of intra and inter H-bonds than WT except for G476S MT, which formed a lesser number of inter H-bonds. Further, secondary structures such as β-Sheets, helices, β-bridges, bends, turns and coils play an essential role in providing distinct shape and pattern to the protein structures. Each moiety (Sheet, helix, bridge, bend, turn and coil) was further elucidated independently using a dictionary of secondary structure of protein (DSSP) approach and found that helix and β-bridge contents were increased and decreased in MT and WT, respectively. Moreover, all secondary structure moieties remain altered in both WT and MT proteins (Fig. S5).

### 3.4 Essential motions of WT and MTs

Essential dynamics which exploited PCA was conducted to elucidate the collective or fundamental motions of WT and MTs. Collective movements are helping to study the conformational changes that existed in the protein, thus reflecting the biological function of that protein. We used stable MD simulation trajectories (125-150ns) to measure the protein dynamics and sketched the first 20 eigenvectors with their corresponding eigenvalues. Out of which, the first 3 eigenvectors accompanied with maximum motions and their cumulative percentage were 77.04, 87.4, 65.27, 67.33, 63.26 and 68.62% for WT, V367F, R408I, G476S, V483A, and N501Y MTs, respectively (Fig. 4A & B). Moreover, we found a similar pattern of fluctuations in PC1, PC2, and PC3 as we found during RMSF analysis (Fig. S6). To ensure the biological relevant motions of PCs, we calculated cosine content. We found that the first three PCs of both WT and MT proteins exhibited a low value of cosine (≤0.2), thus indicating that motions were not due to random diffusions (Table 1). Conformational sampling of WT and MTs were enquired by plotting and comparing the projection of the first three PCs in phase space in a manner such as PC1 vs PC2, PC1 vs PC3 and PC2 vs PC3. During the projection of PC1 vs PC2 analysis, N501Y occupied a larger sub space whereas the rest of mMTs and WT engaged smaller subspaces (Fig. 4C & D). Similar patterns were also observed when projecting the PC1 vs PC3 and PC2 vs PC3 in phase space (Fig. S7). The above results showed that N501Y mutant undergoes large conformational changes. The conformational changes at the structural level were further elucidated by sequentially superimposing the 30 frames of first PC for all WT and MTs (Fig. 5). The variations in the motions were mostly found in extended loop regions of MTs as well as WT. Moreover, N501Y MT exhibited additional motions near N-terminal besides the motion at extended loop region (Fig. 5F). Furthermore, the direction and strength of motions were also examined through plotting the porcupine plot of respective PCs (Fig. 5). In porcupine structure, the arrows and length of cone represented direction and magnitude of motions. Different types of motions were exhibited by the extended loop of WT and MT proteins. The upward motions were found in the extended loop of WT and V483A MT (Fig. 5A & E), while downward motions have existed in the loop regions of V367F, R408I and G476S MTs (Fig. 5B-D). Moreover, mixed motions were found in N501Y MT, as downward and upward motions exhibited by the extended loop and rotational motions were found in N-terminal region (Fig. 5F). ED results showed that N501Y MT exhibited a large conformational change at the extended loop and other random moieties of protein, implying that these regions might play a crucial role in acquiring the stable conformation.

**Table 1.**
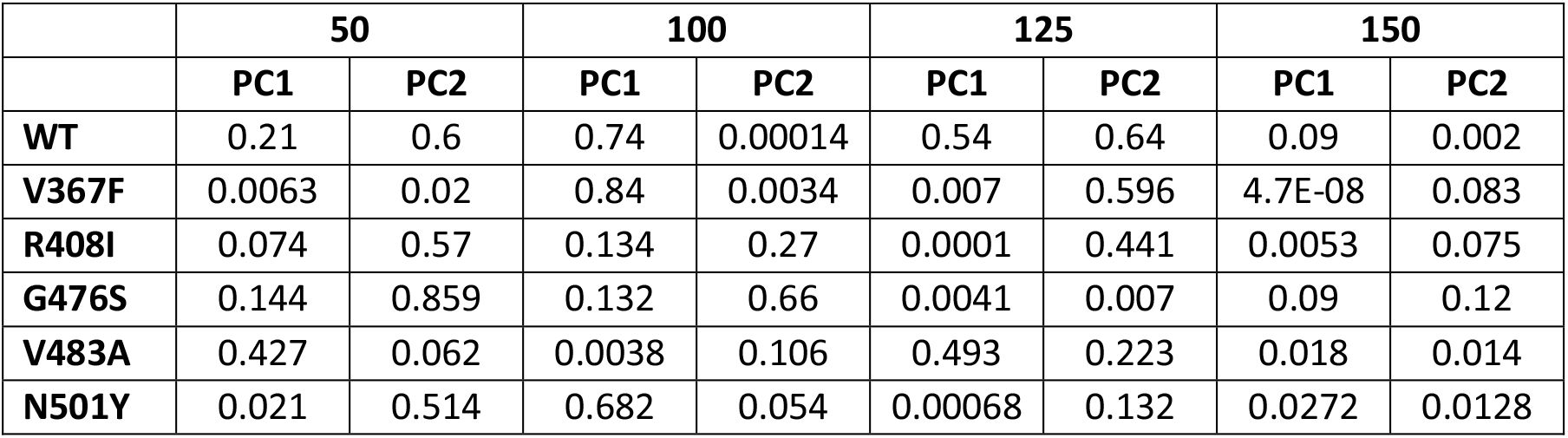
Cosine content analysis of first two principle components (PC1 and PC2)

|  | 50 |  | 100 |  | 125 |  | 150 |  |
| --- | --- | --- | --- | --- | --- | --- | --- | --- |
|  | PC1 | PC2 | PC1 | PC2 | PC1 | PC2 | PC1 | PC2 |
| WT | 0.21 | 0.6 | 0.74 | 0.00014 | 0.54 | 0.64 | 0.09 | 0.002 |
| V367F | 0.0063 | 0.02 | 0.84 | 0.0034 | 0.007 | 0.596 | 4.7E-08 | 0.083 |
| R408I | 0.074 | 0.57 | 0.134 | 0.27 | 0.0001 | 0.441 | 0.0053 | 0.075 |
| G476S | 0.144 | 0.859 | 0.132 | 0.66 | 0.0041 | 0.007 | 0.09 | 0.12 |
| V483A | 0.427 | 0.062 | 0.0038 | 0.106 | 0.493 | 0.223 | 0.018 | 0.014 |
| N501Y | 0.021 | 0.514 | 0.682 | 0.054 | 0.00068 | 0.132 | 0.0272 | 0.0128 |

**Fig. 4.**
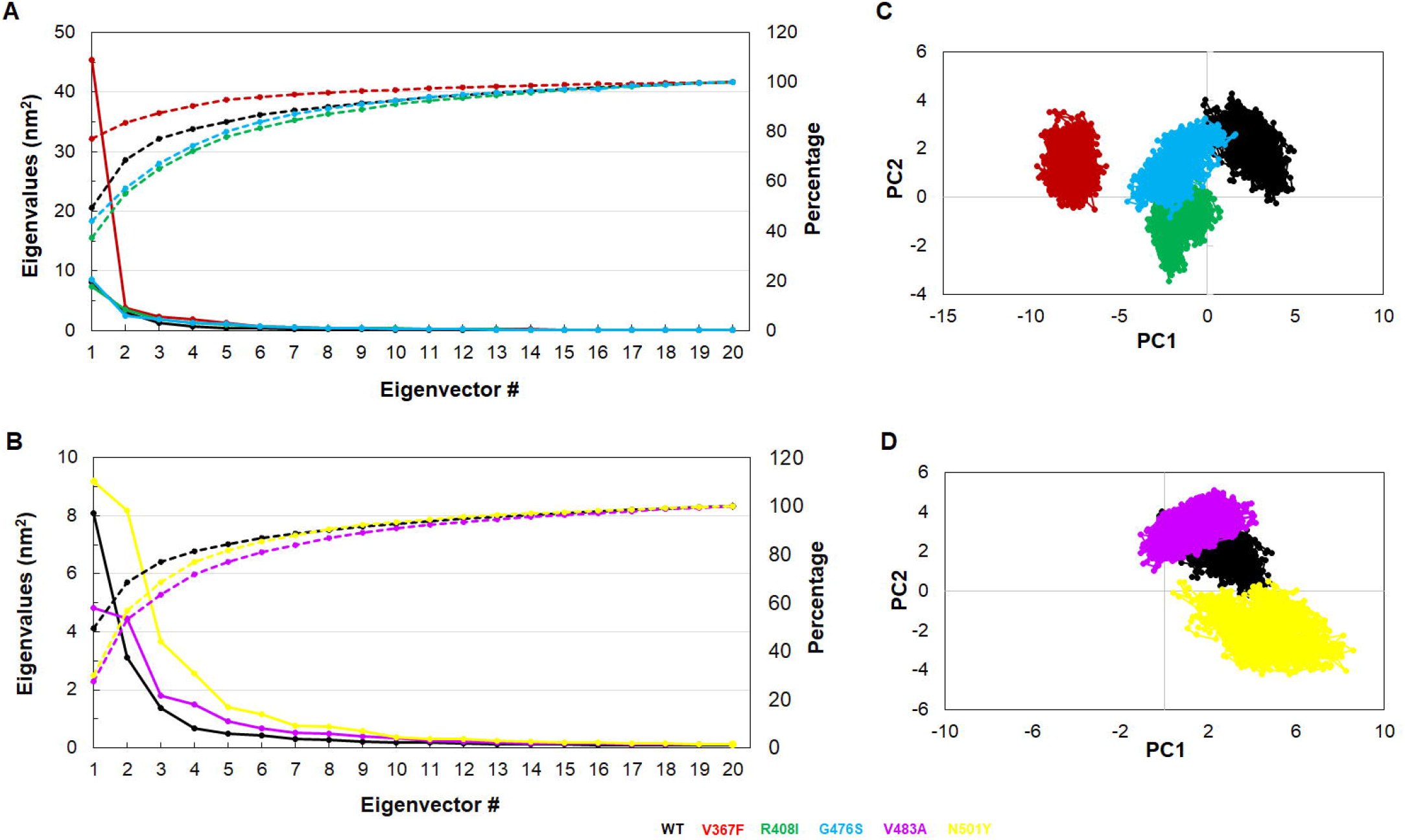
Essential dynamics analysis. (A) Plot of first 20 eigenvectors with associated eigenvalues (solid lines) and their cumulative percentage (dotted lines) of WT, V367F, R408I and G476S MTs, (B) Plot of first 20 eigenvectors with associated eigenvalues (solid lines) and their cumulative percentage (dotted lines) of WT, V483A and N501Y MTs, (C) Projection of PC1 vs PC2 of WT, V367F, R408I and G476S MTs and (D) Projection of PC1 vs PC2 of WT, V483A and N501Y MTs. WT, V367F, R408I, G476S, V483A and N501Y MTs were labelled in black, red, green, blue, magenta and yellow colour, respectively.

**Fig. 5.**
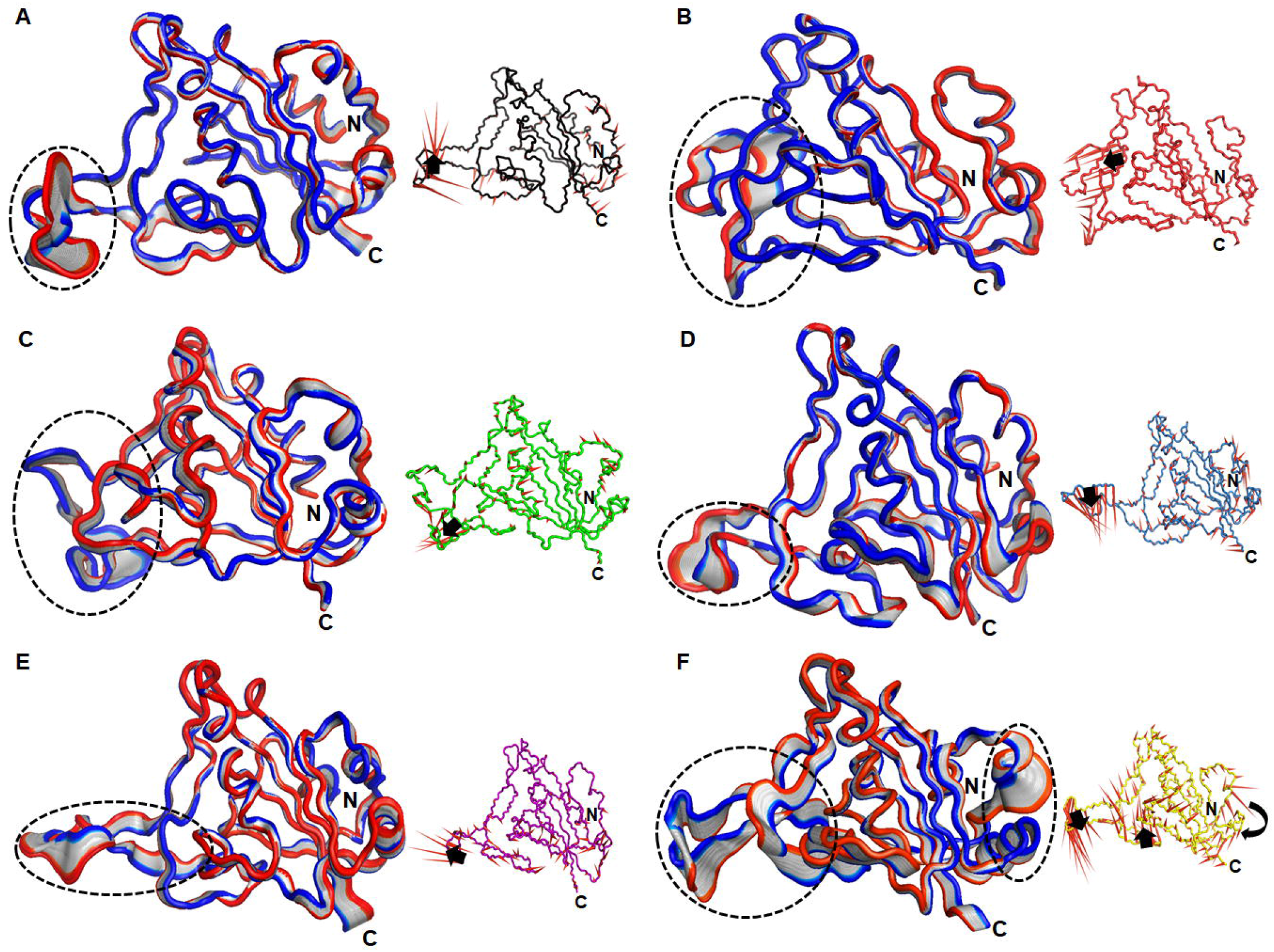
Major motions analyses of first PC. 30 frames of PC1 were sequentially superimposed and associated motions were depicted in porcupine structures of (A) WT, (B) V367F, (C) R408I, (D) G476S, (E) V483A and (F) N501Y. Intensities and nature of motions were highlighted in dotted circles and arrows in the protein 3D and porcupine structures, respectively.

### 3.5 WT and MTs showed almost similar residues interaction networks

To explore the essential residues involved in the interaction and signalling transduction in the protein, residues interaction network (RIN) analysis was performed for all MTs and WT. MD optimized 3D structures were employed to calculate the centrality and to decipher its connectivity. Here, we computed betweenness centrality (C_B_), which is a critical centrality based on the shortest route between the nodes and considered the residues with a threshold C_B_ value ≥0.2 (Fig. 6). WT and other mutants such as V367F, R408I, G476S, and V483A showed only one residue (Tyrosine453) as par with threshold C_B_ value which is located in the β-sheet region of the protein (Fig. 6A-E). In case of N501Y, four residues, namely Valine 350, Tyrosine451, Tyrosine453 and Tyrosine495 (V350, Y451, Y453 and Y495) were had higher C_B_ values (Fig. 6F). Interestingly, all residues except Y453 lies at the loop region of protein which indeed indicating an important region in the residual signalling of the protein.

**Fig. 6.**
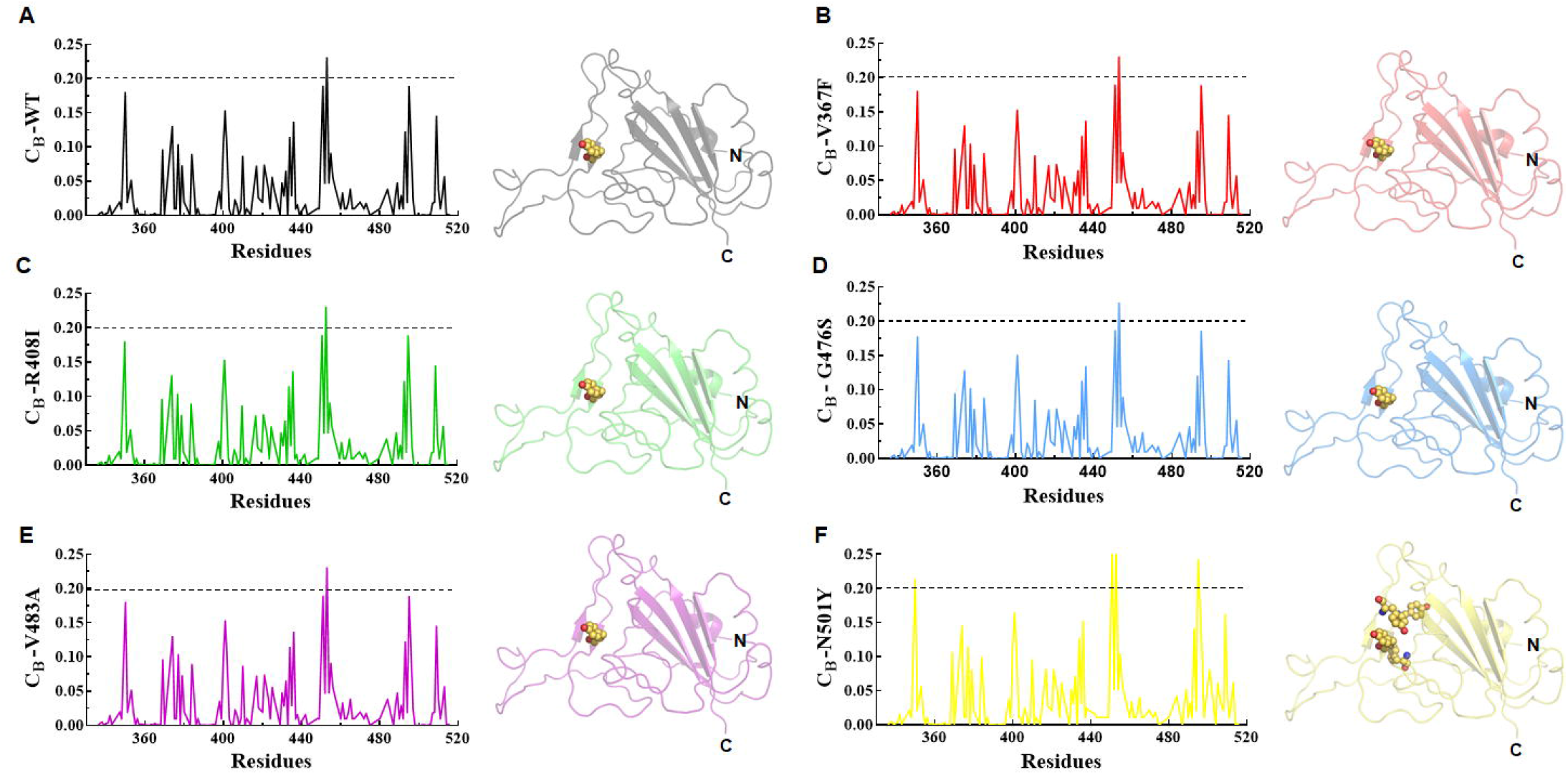
Network centrality analysis. Betweenness centrality (C_B_) values of (A) WT, (B) V367F, (C) R408I, (D) G476S, (E) V483A and (F) N501Y. Black dotted line represented the cut off value used to filter the important residues. Right panel of each graph denoted 3D structure displayed in cartoon modes in which important residues shown in sphere.

### 3.6 MTs had high binding affinities than WT

Protein–protein (p-p) interaction is a physical association of two or more proteins due to biochemical events driven by many biological forces like Van der Waals, electrostatics and polar interactions. It plays a decisive role in the overall physiology of the cell and its biological process, including signal transduction. Structure based molecular docking approach was widely used to predict the orientation of one protein with respect to other and the key residues involved in the interaction (Maurya et al., 2020). Here, we employed the HADDOCK server to study the p-p docking of ACE2 (angiotensin-converting enzyme 2) receptor and SpikeS1 RBD of WT and MTs. HADDOCK predicted different clusters for each WT- and MT-protein complexes out of which cluster accompanied larger structures had the lowest HADDOCK score, which indicated the p-p complex of high binding strength. Topmost clusters size of ACE2-WT, -V367F, -R408I, -G476S, -V483A and -N501Y MT complexes were had 168, 169, 127, 175, 176 and 62 structures, respectively (Table 2). HADDOCK score of topmost clusters for ACE2-WT, -V367F, -R408I, -G476S, -V483A and -N501Y MT complexes were −122.9 (±3.3), −122 (±2.2), −114.2 (±2), −121.4 (±2.6), −122.1 (±2.3) and −124.4 (±3.1), respectively (Fig. 7; Table 2). Various energies such as van der Waals, electrostatic, desolvation and restraints violation energies significantly contributed to determining the binding affinities of all WT and MTs p-p interactions (Figs. S8&S9). During binding energy analysis, electrostatic energy had a major contribution in total binding energy which were −212.3 (±27.8), −185.3 (±29.3), −213.5 (±14.2), −244.6 (±9.7), −214.6 (±18.2) and −210.4 (±20.3) for ACE2-WT, -V367F, -R408I, -G476S, -V483A and -N501Y MT complexes, respectively (Table 2). Additionally, all p-p complexes were stabilised by the ample amount of hydrophilic as well as hydrophobic interactions (Figs. S10&S11; Table S6). Furthermore, p-p binding free energies were also verified through the change in binding affinities of ACE2-WT and -MT complexes by using mCSM-PPI server and we found that N501Y MT exhibited a higher value of affinity changes (ΔΔG= −2.147 Kcal/mol) as compared to WT and other MTs (Table S7). The above results indicated that N501Y MT had a high binding affinity to the host ACE2 protein, V367F and V483A have similar binding affinities to that of WT, while R408I and G476S have lower binding affinity to an ACE1 receptor protein.

**Table 2.**
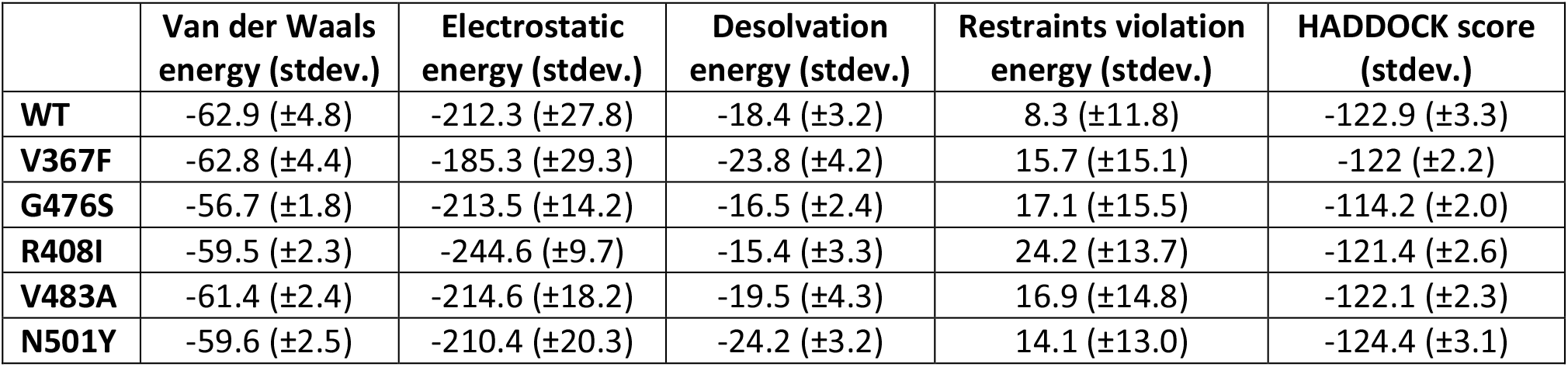
Protein-protein (ACE2-RBD) interaction energies analyses.

|  | Van der Waals<br>energy (stdev.) | Electrostatic<br>energy (stdev.) | Desolvation<br>energy (stdev.) | Restraints violation<br>energy (stdev.) | HADDOCK score<br>(stdev.) |
| --- | --- | --- | --- | --- | --- |
| **WT** | -62.9 (±4.8) | -212.3 (±27.8) | -18.4 (±3.2) | 8.3 (±11.8) | -122.9 (±3.3) |
| **V367F** | -62.8 (±4.4) | -185.3 (±29.3) | -23.8 (±4.2) | 15.7 (±15.1) | -122 (±2.2) |
| **G476S** | -56.7 (±1.8) | -213.5 (±14.2) | -16.5 (±2.4) | 17.1 (±15.5) | -114.2 (±2.0) |
| **R408I** | -59.5 (±2.3) | -244.6 (±9.7) | -15.4 (±3.3) | 24.2 (±13.7) | -121.4 (±2.6) |
| **V483A** | -61.4 (±2.4) | -214.6 (±18.2) | -19.5 (±4.3) | 16.9 (±14.8) | -122.1 (±2.3) |
| **N501Y** | -59.6 (±2.5) | -210.4 (±20.3) | -24.2 (±3.2) | 14.1 (±13.0) | -124.4 (±3.1) |

**Fig. 7.**
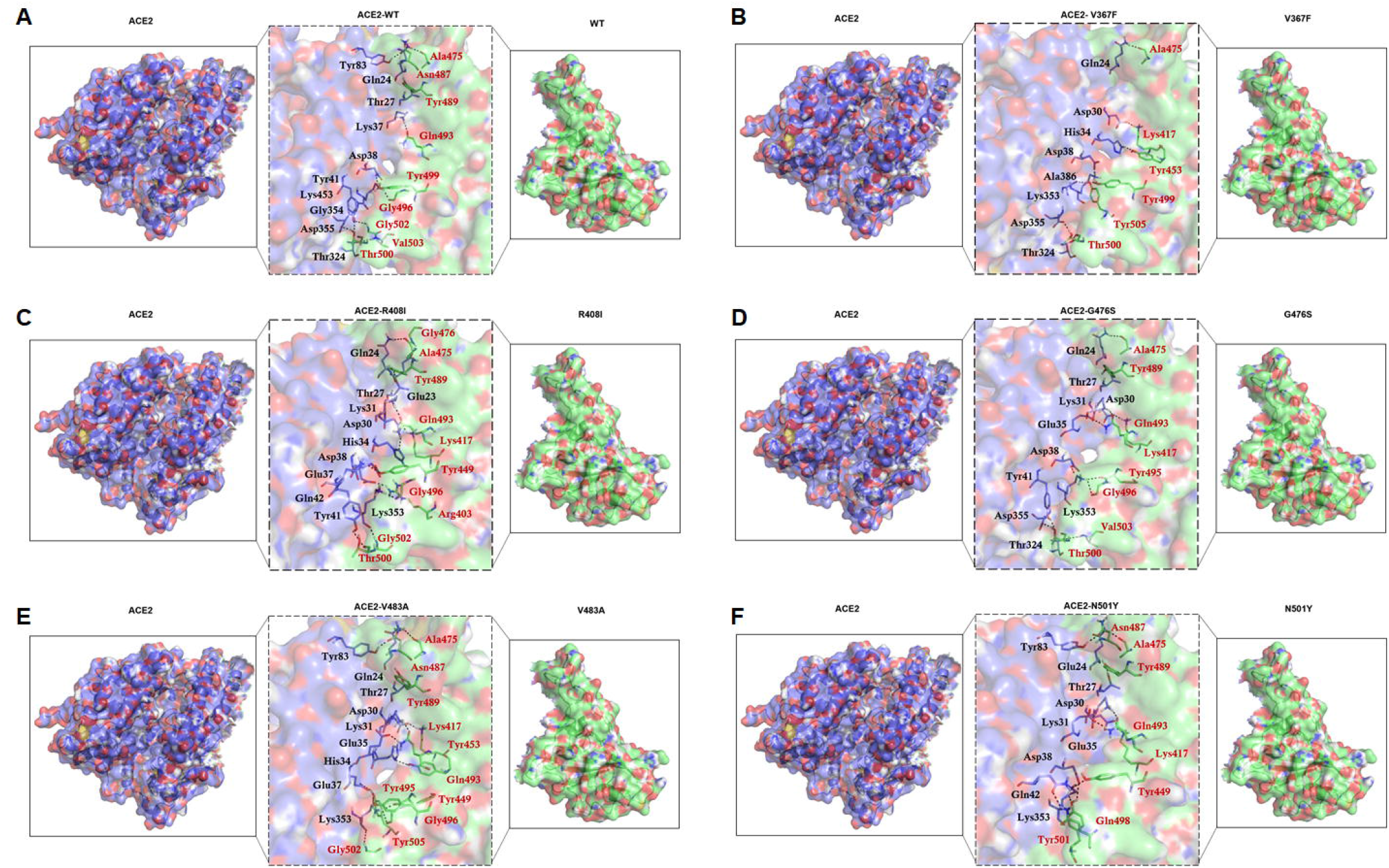
Protein-protein (ACE2-SpikeS1 RBD) interaction study. (A) ACE2-WT complex, (B) ACE2-V367F complex, (C) ACE2-R408I complex, (D) ACE2-G476S complex, (E) ACE2-V483A complex and (F) ACE2-N501Y complex. ACE2 and SpikeS1 RBD were shown in blue and green surface cartoon modes, respectively. Interacting residues of ACE2 (Black) and SpikeS1 (Red) RBD were represented as stick modes with 3 letter amino acid code.

## 4. DISCUSSION

Viruses, particularly RNA viruses have a marvellous tendency to mutate themselves as compare with DNA viruses, resulting in the rapid emergence of new strains (Elena & Sanjuan, 2005). There are currently thousands of SARS-CoV2 variants reported in which the speed and intensity of spreading some of them (variants) are explained, more than by their genetic differences, by the habits of population, and the effectiveness of epidemiological surveillance policies (Hodcroft et al., 2020). SARS-CoV2 is a novel epidemic strain of *Betacoronavirus* responsible for the widespread viral pandemic COVID-19 (Wong & Saier, 2021). The virus gets into the cell with the help of spike glycoprotein (S1) present on the surface of the virus envelope which interacts with the ACE2 receptor of host cell (Korber et al., 2020). Receptor binding domain (RBD) of SpikeS1 protein interacts with ACE2 receptors and are more prone to mutations resulting in the formation of new strains and also help the viruses to escape from different therapies (Lan et al., 2020). The RBD consists of a core region of β-sheets and an extended loop region known as receptor binding motifs that provide a binding surface to the ACE2 protein (Yan et al., 2020). Before developing a specific and effective drug, one must understand the molecular interaction of ACE2 and SpikeS1 protein of different strains. Therefore, the present study has critically analysed the effect of various mutations on the structural stability and dynamics of RBD of SpikeS1 protein by manifesting different MTs or strains of SARS-CoV2 and assessed the binding affinities with host (human) ACE2 receptor.

The tertiary structure of ACE2-SpikeS1 RBD complex was taken from PDB and used as a template for creating various SARS-CoV2 mutants. Qualities of generated mutant models were verified by inspecting the stereochemical and geometrical properties through different model validation tools and found that all MTs (V367F, R408I, G476S, V483A and N501Y) had good profiles of Ramachandran plots, acceptable scores of ProSA and QMEAN, similar to WT protein, implying that all MTs along with WT had better stereochemical and geometrical properties. To explore the structural and functional properties, the optimised RBD MTs were subjected to MD simulations of 150ns simulation time. MD simulation is a promising technique to study the stability and dynamics of macromolecules at an atomic level in a realistic manner, which is hard to achieve through experimental methods (Kumar et al., 2017; Kumar et al., 2020). Different parameters such as RMSD, RMSF and Rg were measured to monitor the stability and dynamics of WT and MT proteins. RMSD results suggested that WT as well as MT proteins displayed consistent behaviour with higher RMSDs were found in V367F, R408I and V483A MTs as compared to WT, G476S and N501Y MTs. RMSF analysis was performed to study the residue and structural level alterations in WT and MTs and found residues like Gly476, Ser477, Thr478, Asn481, Gly482 and Val483 were highly fluctuated and located at loop regions of almost all MTs and WT. Loops are essential moieties of protein that are highly mobile and act as a crucial site for ligand interactions and help in acquiring the stable conformation of proteins (Baiesi et al., 2019). Hence, the mobility of residues in the loop regions in WT and MTs help in the interactions of RBD to partner protein such as ACE2.

Next, we have analysed the structural properties of all MTs along with WT by measuring SASA, H-bonds and secondary structure units. The core region of protein mainly consists of hydrophobic residues that act as a driving force during protein folding. The solvent interaction intensity of amino acid and the core of protein is proportional to the total surface area exposed to these environments (Kumar & Saran, 2021). Therefore, SASA of WT and MTs were measured to monitor the surface area available for the surrounding environment and found that all MTs displayed higher SASA then WT and hydrophobic SASA contributed slightly more in comparison to hydrophilic SASA as RBD of protein mostly consist of core followed by loop regions. Hydrogen bonding (H-bond) is an important parameter for providing stability and strength of protein-ligand interaction properties of a particular protein. The stability and protein-ligand interaction capacity were measured by calculating the intra (protein-protein) and inter (protein-solvent) H-bonds (Pace et al., 2014). Our results suggested that almost all MTs formed a high number of intra and inter H-bonds compared to WT, demonstrating that MTs formed highly stable protein structures and have a higher tendency of ligand interactions. Furthermore, secondary structure analyses of all MTs and WT indicated that alterations in the random moieties of protein compromised with changing in α-helix and β-sheet contents.

Essential dynamics (ED), a typical application of PCA, which is used to extract the biological relevant motions by utilising the MD simulation trajectories (Berendsen & Hayward, 2000). Equilibrated MD trajectories were harnessed for constructing the covariance matrix and series of eigenvectors with associated eigenvalues were plotted for all MTs and WT. The first three eigenvectors were used to study the conformational subspace of MTs and WT protein as they are mostly encompassing the major motions. ED results suggested that, except for N501Y MT, all other MTs were had similar conformational subspace as with WT, while N501Y MT covered larger conformational space. Moreover, the structure level inspection from the first eigenvector (PC1) revealed that the higher motions were accompanied by extended loop region of the protein. The biological function of a protein is characterizing by constituent residues and their chemical properties. These chemical versatilities of amino acids determine the overall topology of protein and its behaviour in the surrounding environment (Przulj et al., 2004). Different residues in the protein interact and help in the flow of signal, which is better elucidated by the residues interaction network (RIN). In this analysis, an entire protein considered as a network in which residues and interactions are plotted in the form of nodes and edges, respectively (Davis et al., 2015). Three centralities were computed out of which betweenness centrality (C_B_) is most important concerning to protein function (Kumar et al., 2020). RIN results suggested that Tyrosine453 was an important residue in WT and V367F, R408I, G476S, and V483A MTs, while more than one amino acid had role in signalling in N501Y MT. The above residues are mainly located on or near the 5^th^ and 6^th^ β-sheets of RBD, demonstrating that this region might be crucial for the functioning of WT and MT proteins.

SARS-CoV2 invades host cells through the interaction of RBD of SpikeS1 unit to the host ACE2 receptor. Qualitatively, the infectivity of diverse SARS-CoV2 strains in the host is proportional to the binding affinity of SARS-CoV2 SpikeS1 RBD of each strain to the ACE2 receptor of host cells (Mittal et al., 2020). Therefore, the assessment of binding affinity of different MTs to host cells is vital for understanding the infectiousness of SARS-CoV2. The binding affinity of WT and MTs RBD to ACE2 was elucidated by the HADDOCK protein-protein (p-p) docking tool. P-p docking results suggested that V367F and V483A MTs have similar binding affinities to WT, while R408I and G476S had fewer binding affinities. Moreover, N501Y MT had a higher binding affinity than the rest of MT and WT proteins. Binding affinity analysis indicated that N501Y MT binds to host ACE2 protein with maximum binding strength, thus contributing highly infectious activity, as also reported previously (Luan et al., 2021).

Proteins are having a dynamic personality and are central to the molecular function of a cell. Any change in amino acid residue led to the changing in overall topology and hence affected their functions. The molecular and functional consequences of change in amino acids are better observed in the current pandemic (COVID-19) as the point mutations in the SpikeS1 RBD of SARS-CoV2 results in the generation of thousands of variants or strains (Chen et al., 2020). All countries of the world are witnessed the infectiousness of these strains. The alteration in amino acids showed both local and the global impact on the topology of protein and in turn, altered the interacting residue, which ultimately affected the binding affinity of two partner proteins (ACE2-SpikeS1 RBD). In V367F MT, a nonpolar hydrophobic amino acid has aliphatic side chain (Valine) is mutated to an aromatic hydrophobic residue (Phenylalanine) that enhanced the structural stability of the spike protein, thus facilitating more efficient binding to the host ACE2 receptor. In case of R408I MT, there is change in overall charge of protein residue (Arginine), which is mutated to the residues having a neutral charge (Isoleucine). This also affects overall topology of protein as mutated residue have a smaller size than WT residue, resulting in the decreasing of overall binding affinity as reported during p-p interaction. In G476S MT, Glycine of WT residues is mutated to Serine which affects the overall flexibility of protein as Glycine is more flexible than Serine. This led to a decrease in the stability and binding affinity of G476S MT, as noticed during MD simulation and p-p docking. In V483A MT Valine, a large hydrophobic protein residue is mutated to Alanine, a small size hydrophobic residue that alters the overall dimension of the protein, thus help in increasing the binding affinity. And, in case of N501Y MT, a substitution of Asparagine to Tyrosine was found in the binding motif of SpikeS1 RBD, which enhances the binding affinity to its protein partner and ultimately leads to a stable virus-host relationship. Due to strong binding, this mutation also assists the virus in evading antibody neutralization.

The present scenario of global pandemic, where the world keeps an eye towards the vaccine at utmost level. In this regard, the computational approach has the potential to unravel the mechanistic behaviour of RBD MTs and its interaction with ACE receptor at an atomic level. However, certain limitations are associated with the current computational study, which may be overcome by experimental methods. Therefore, experimental methods are highly urged to support or validate the findings of the current research. Nevertheless, our study provides computational evidence of the strength of highly infectious strains, which would help in design a more specific drug to combat the current pandemic.

## CONCLUSION

Noteworthy infectivity of severe acute respiratory syndrome-coronavirus 2 (SARS-CoV2) is due to its fast-mutating ability thus led to extended infection. In this investigation, the computational methods such as MDS and p-p docking have been applied and revealed that mutant strains of SARS-CoV2 formed stabled protein structures and had higher binding affinities to host protein. Hence, the present study provides the molecular evidence of this endless pandemic which would help in designing a more specific inhibitor.

## ACKNOWLEDGEMENTS

Rakesh and Rahul thank Indian Council of Medical Research and University Grant Commission, respectively for financial support. H.G. acknowledged DST (Department of science and technology), India for financial assistance.

## AUTHOR CONTRIBUTION

Rakesh K. and P.T. conceptualised and designed the study. Rakesh K. and Rahul K. performed experiments and analysed the data. H.G. help in literature study. Rakesh K., Rahul K. and H.G. wrote the manuscript. P.T. provided laboratory infrastructure. All authors read and approved the final version of manuscript.

## DATA AVAILABILITY STATEMENT

All datasets generated during this study are included in the article/supplementary material.

## List of abbreviations

ACE2: Angiotensin converting enzyme2
COVID-19: Coronavirus disease-2019
DSSP: Dictionary of secondary structure of protein
ED: Essential dynamics
GROMACS: Groningen machine for chemical simulation
HADDOCK: High ambiguity driven docking
MDS: Molecular dynamics simulation
MT: Mutant
PCA: Principle component analysis
PDB: Protein data bank
ProSA: Protein structure analysis
QMEAN: Qualitative model energy analysis
RBD: Receptor binding domain
Rg: Radius of gyration
RIN: Residues interaction network
RMSD: Root mean square deviation
RMSF: Root mean square fluctuation
SARS-CoV2: Severe acute respiratory syndrome-coronavirus 2
SASA: Solvent accessible surface area
SAVES: Structure analysis and verification server
WT: Wildtype

