## Supplemental File for "Computational investigation reveals that the mutant strains of SARS-CoV2 are highly infectious than wildtype"

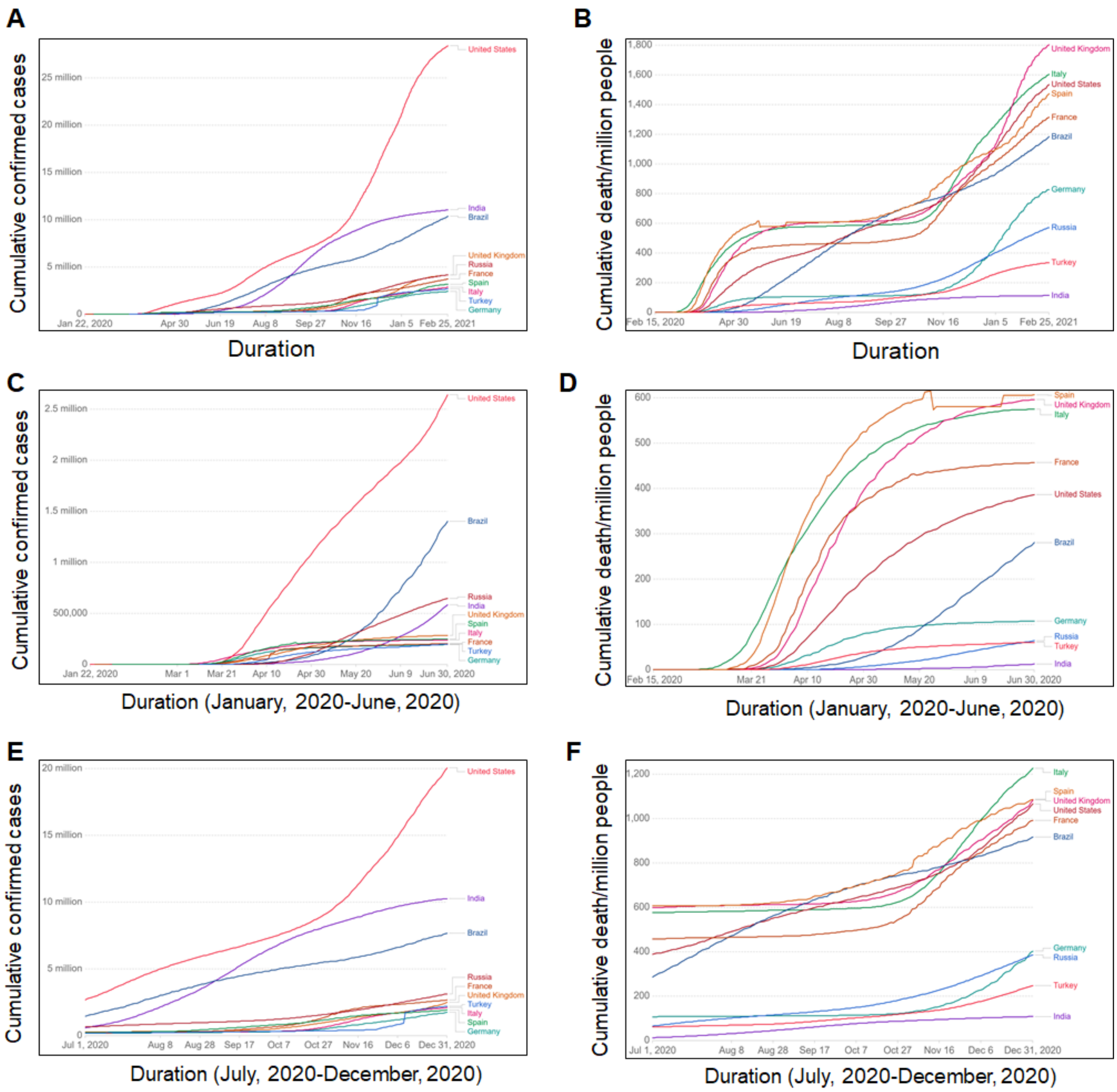


**Fig. S1** **Coronavirus pandemic country profile** (till February, 2021)**.** (A) Total conformed cases, (B) Total death/million people, (C) Total confirmed cases of first wave (January, 2020- June, 2020), (D) Total death/million people in first wave, (E) Total confirmed cases of second wave (July, 2020- December, 2020) and (F) Total death/million people in second wave. Different countries were labelled in different colour lines. Data was taken from Coronavirus Pandemic (COVID-19) database. (https://ourworldindata.org/coronavirus).

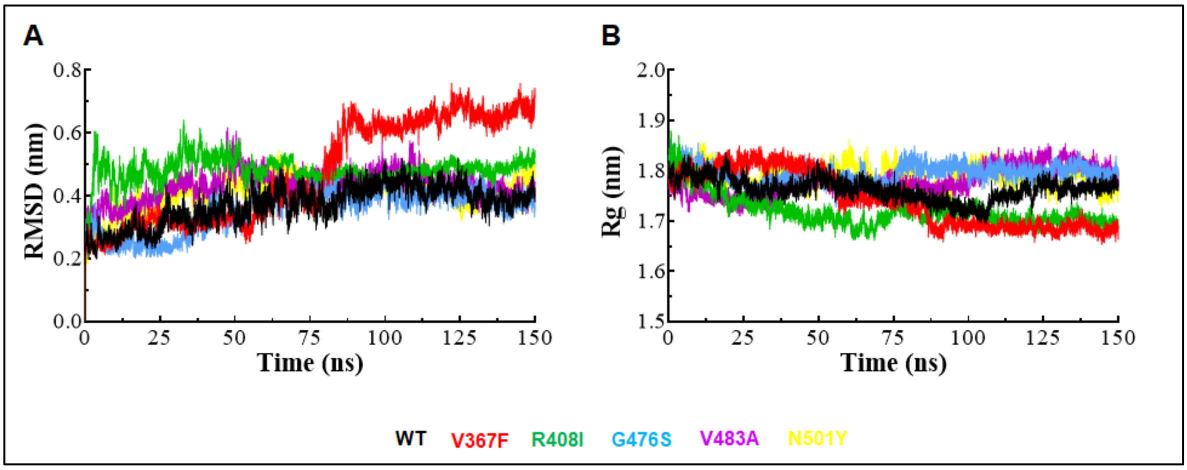


**Fig. S2** **MD simulation profiles of WT and MT proteins.** (A) RMSD plot and (B) Rg plot. WT, V367F, R408I, G476S, V483A and N501Y MTs were labelled in black, red, green, blue, magenta and yellow colour, respectively.

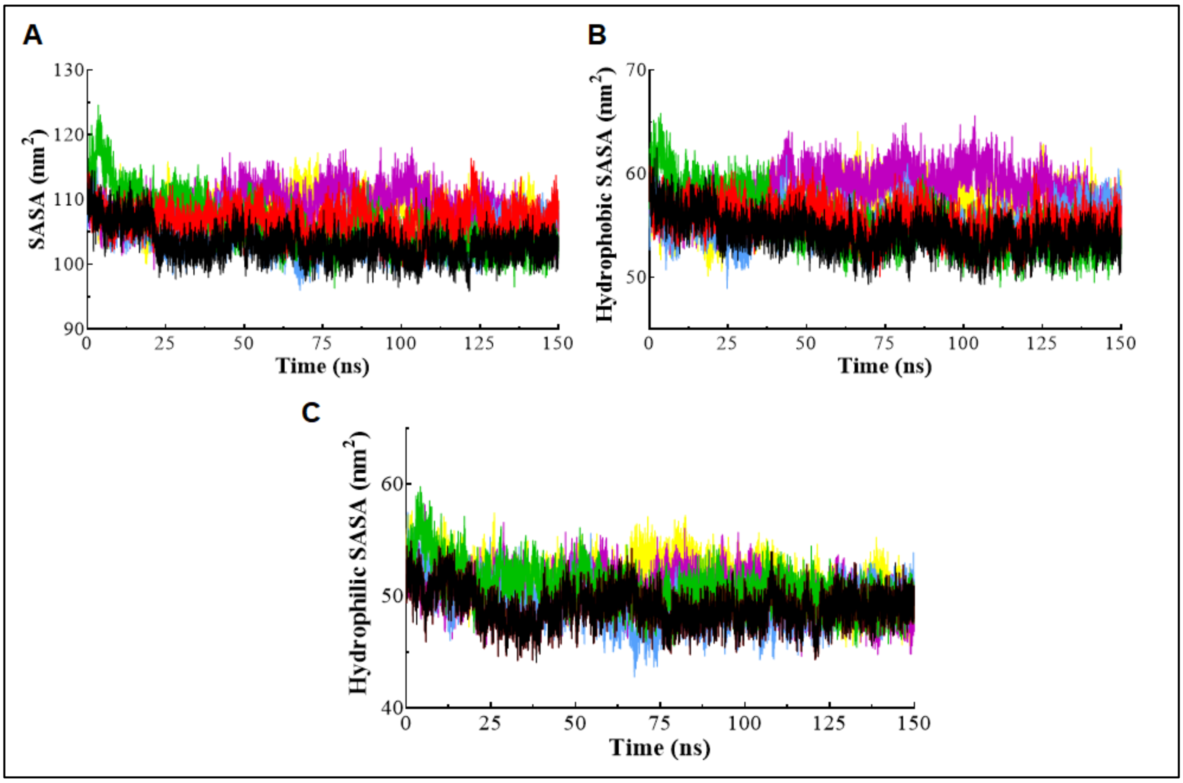


**Fig. S3** **Solvent accessible surface area (SASA) analysis.** (A) Total SASA, (B) Hydrophobic SASA and (C) Hydrophilic SASA. WT, V367F, R408I, G476S, V483A and N501Y MTs were labelled in black, red, green, blue, magenta and yellow colour, respectively.

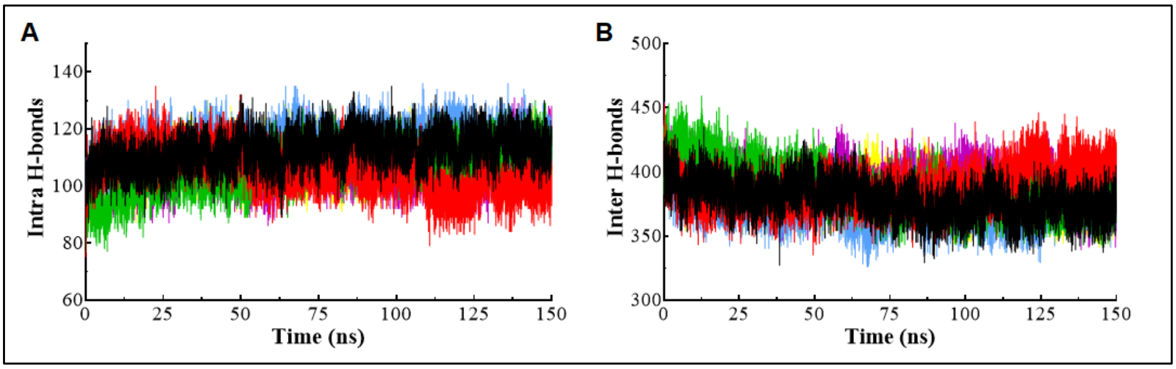


**Fig. S4 Hydrogen bond (H-bond) analysis.** (A) Intra or protein-protein H-bond and (B) Inter or protein-water H-bond. WT, V367F, R408I, G476S, V483A and N501Y MTs were labelled in black, red, green, blue, magenta and yellow colour, respectively.

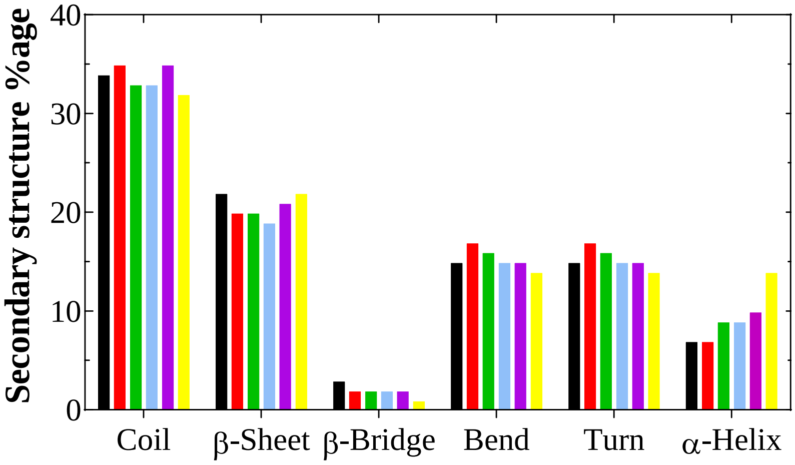


**Fig. S5 Secondary structure analysis of WT and MT proteins.** WT, V367F, R408I, G476S, V483A and N501Y MTs were labelled in black, red, green, blue, magenta and yellow colour, respectively.

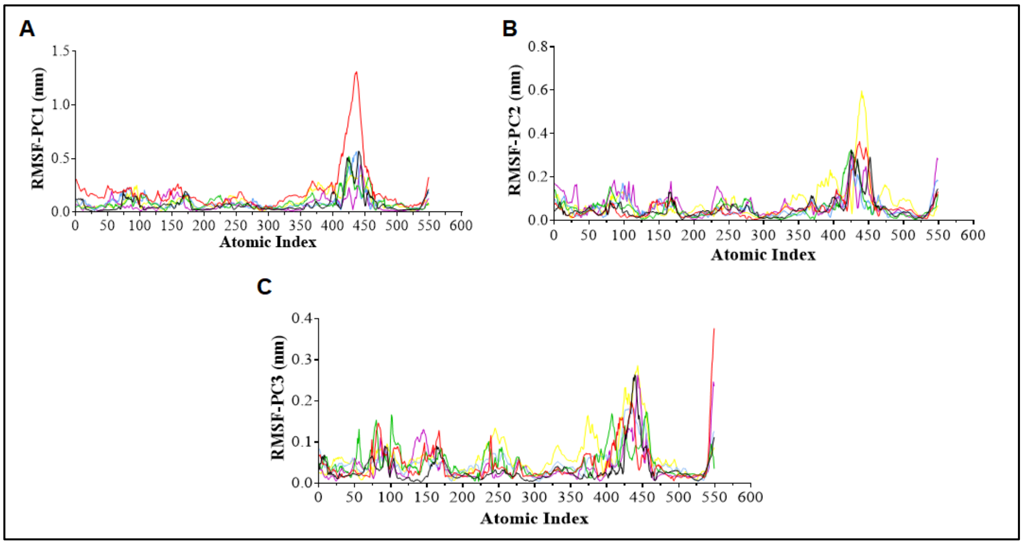


**Fig. S6 RMS fluctuation of first three PCs (PC1, PC2 and PC3) from WT and MTs.** WT, V367F, R408I, G476S, V483A and N501Y MTs were labelled in black, red, green, blue, magenta and yellow colour, respectively.

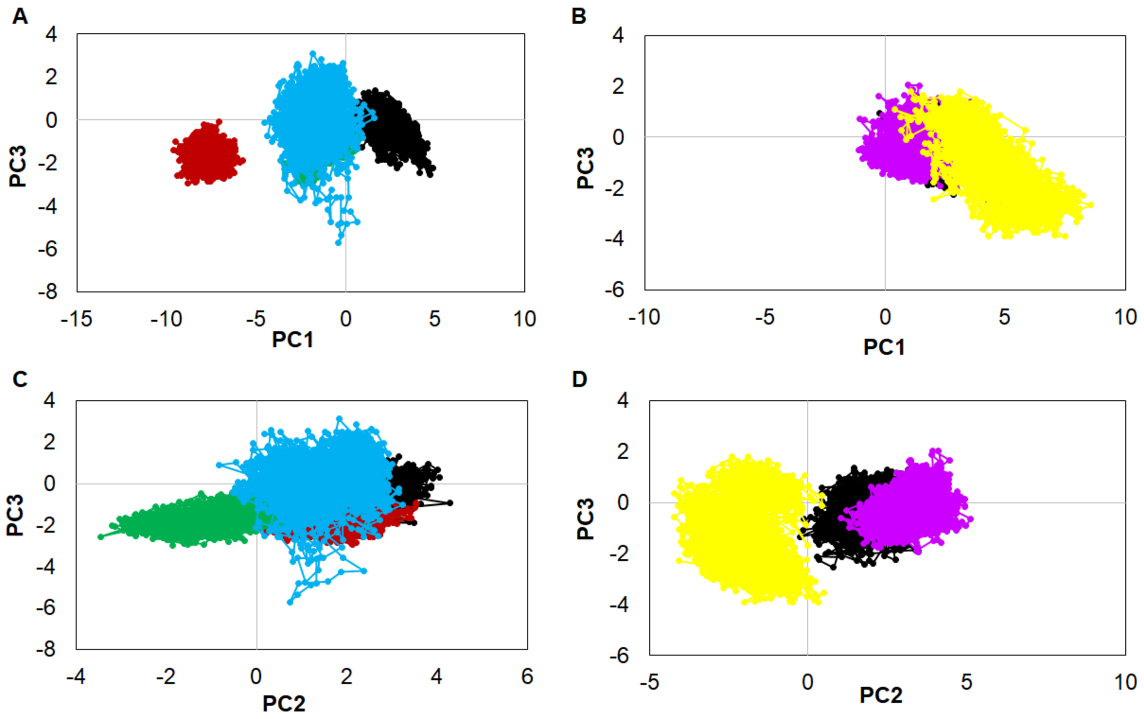


**Fig. S7 Projection of eigenvector1 (PC1) vs eigenvector3 (PC3) and eigenvector2 (PC2) vs eigenvector3 (PC3).** WT, V367F, R408I, G476S, V483A and N501Y MTs were labelled in black, red, green, blue, magenta and yellow colour, respectively.

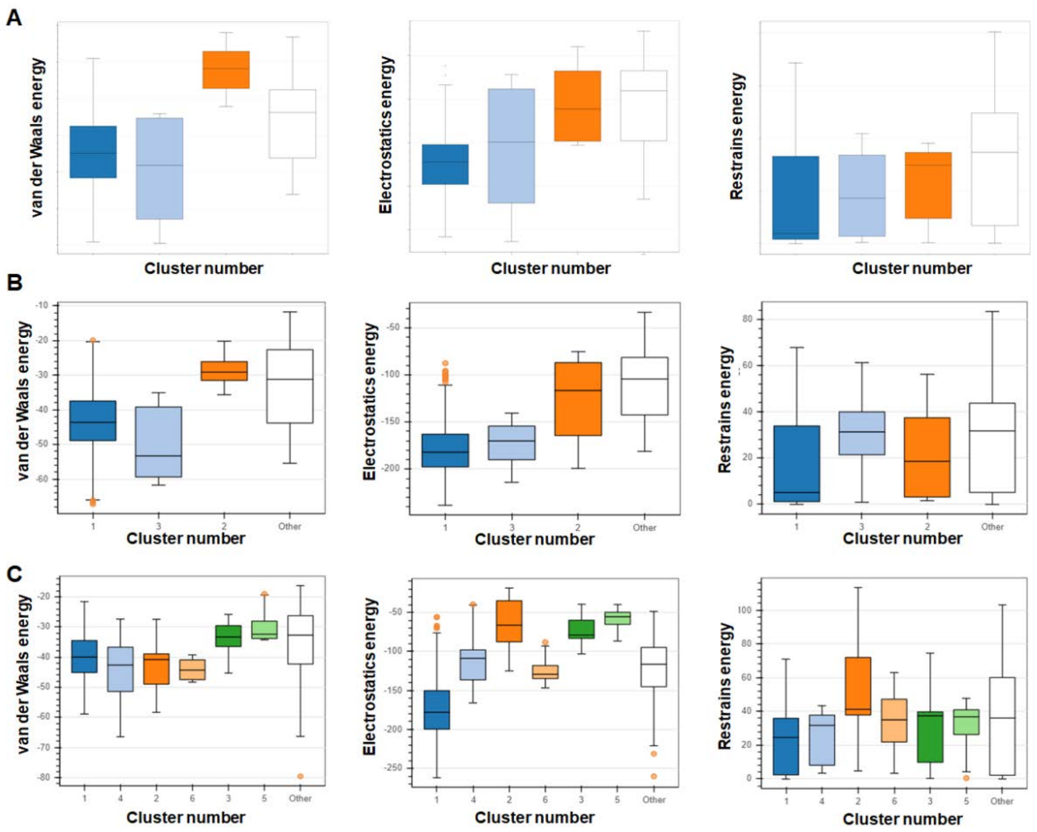


**Fig. S8 Top clusters of protein-protein docking complexes.** (A) ACE2-WT, (B) ACE2-V367F and (D) ACE2-R408I.

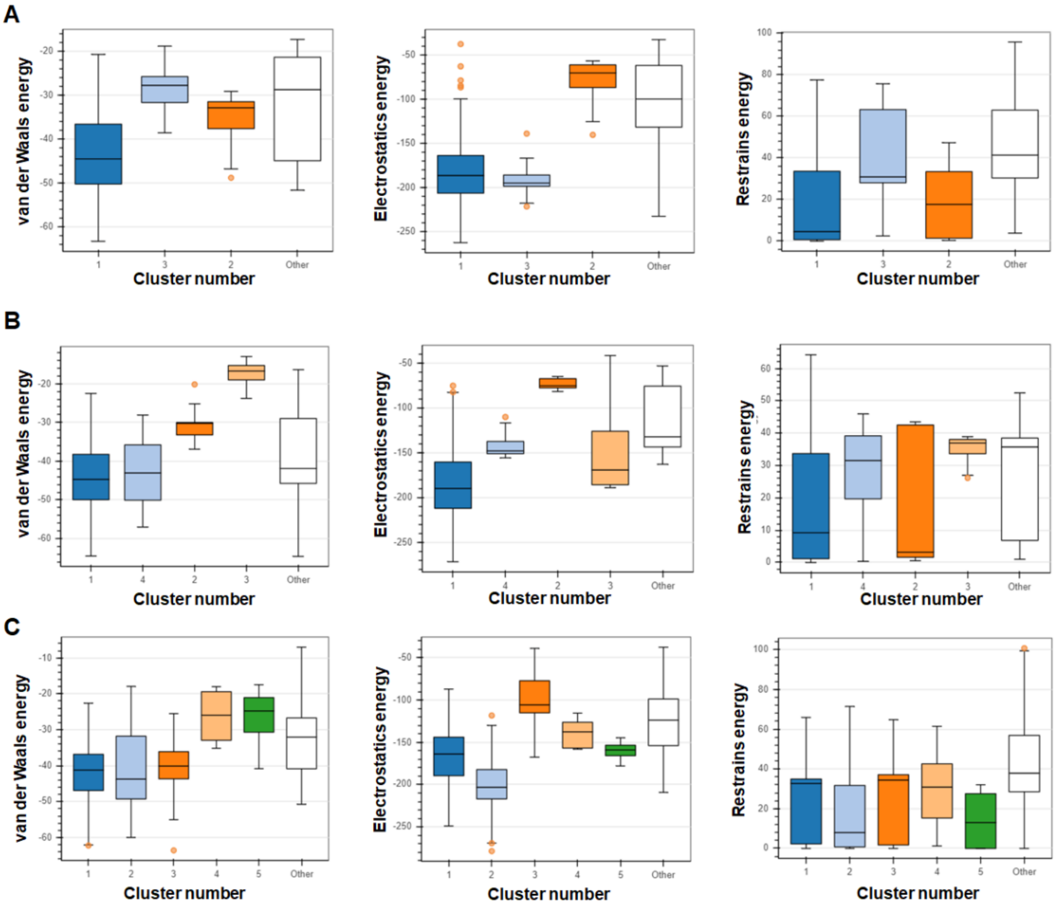


**Fig. S9 Top clusters of protein-protein docking complexes contd**. (A) ACE2-G476S, (B) ACE2-V483A and (C) ACE2-N501Y.

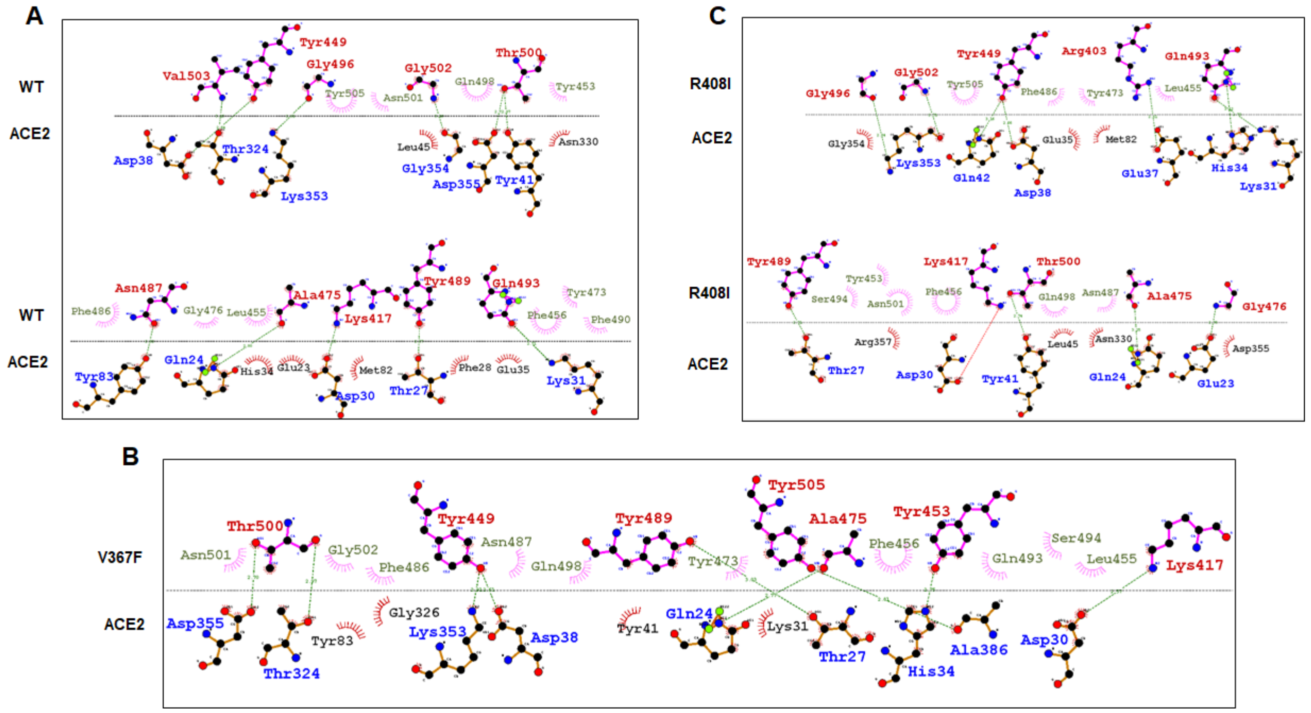


**Fig. S10 2D interaction plots of** (A) ACE2-WT (SpikeS1 RBD), (B) ACE2-V367F and (D) ACE2-R408I complexes. Hydrophobic and hydrophilic residues of ACE2 and SpikeS1 RBD were labelled in Black

(ACE2), light green (SpikeS1 RBD), blue (ACE2) and red (SpikeS1 RBD), respectively. Hydrogen bonds were labelled in green dotted lines.

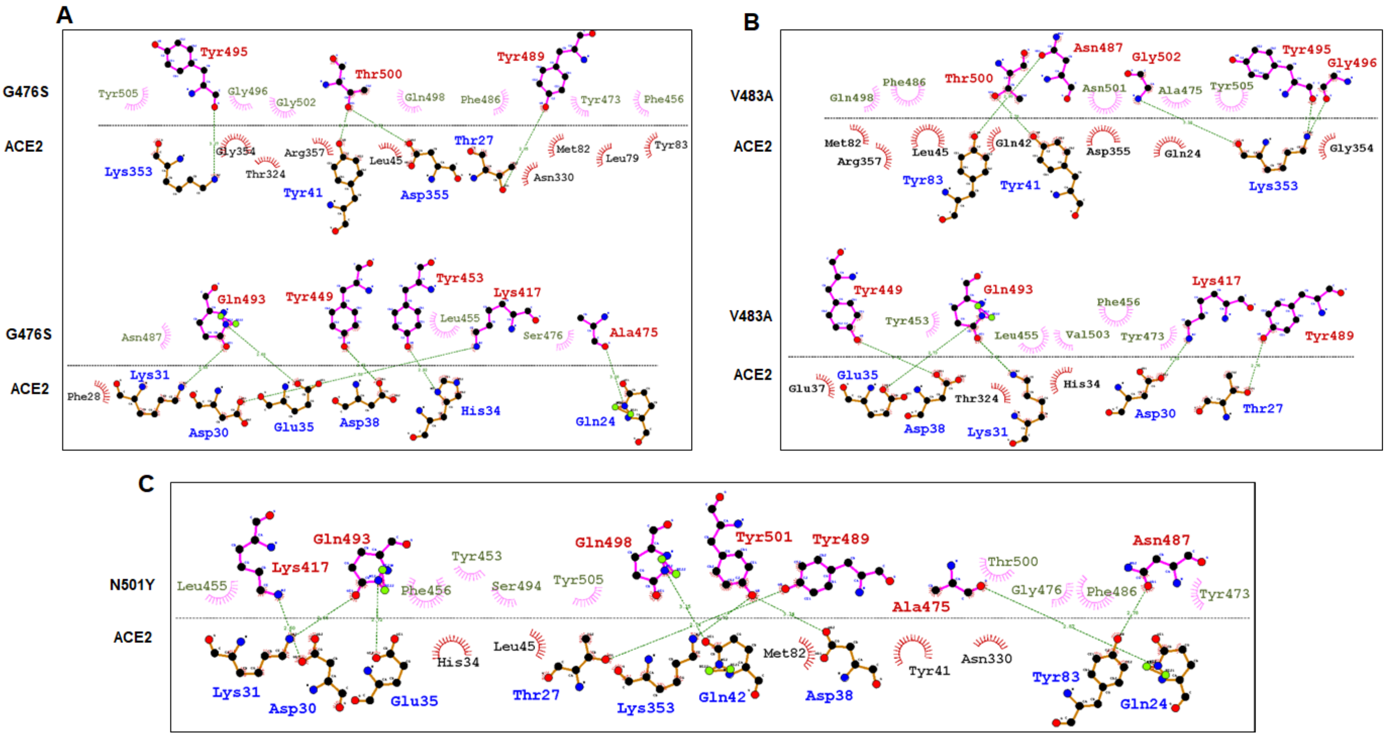


**Fig. S11 2D interaction plots of** (A) ACE2-G476S, (B) ACE2-V483A and (C) ACE2-N501Y complexes. Hydrophobic and hydrophilic residues of ACE2 and SpikeS1 RBD were labelled in black (ACE2), light green (SpikeS1 RBD), blue (ACE2) and red (SpikeS1 RBD), respectively. Hydrogen bonds were labelled in green dotted lines.

### Table S1. Cumulative confirmed COVID-19 cases in million

| Countries | January-June, 2020 | July-December 2020 | January, 2020 -February, 2021 |
|---|---|---|---|
| **USA** | 2.64 | 20.06 | 28.41 |
| **INDIA** | 0.58 | 10.27 | 11.06 |
| **BRAZIL** | 1.40 | 7.68 | 10.39 |

### Table S2. Cumulative confirmed COVID-19 deaths per million

| Countries | January-June, 2020 | July-December, 2020 | January, 2020 -February 2021 |
|---|---|---|---|
| **ITALY** | 575.02 | 1,226.54 | 1,603.89 |
| **USA** | 386.56 | 1,055.29 | 1,535.66 |
| **UK** | 596.28 | 1,084.49 | 1,801.49 |

### Table S3. Structure validation of different MT models along with WT

| Variants | Favoured | Allowed | Disallowed | ProSA (Z-score) | QMEAN (Z-score) |
|---|---|---|---|---|---|
| **WT** | 68.50% | 29.50% | 2% | -5.25 | -7.94 |
| **V367F** | 68.50% | 29.50% | 2% | -5.27 | -7.88 |
| **R408I** | 68.50% | 29.50% | 2% | -5.15 | -8.15 |
| **G476S** | 68.50% | 29.50% | 2% | -5.23 | -7.33 |
| **V483A** | 68.50% | 29.50% | 2% | -5.4 | -7.95 |
| **N501Y** | 68.50% | 29.50% | 2% | -5.8 | -7.99 |

### Table S4. Protein stability prediction through MUpro server

| | | Support vector Machine | | | Neural Network | |
|---|---|---|---|---|---|---|
| **Variants** | **DDG** | **Stability** | **Confidence score** | **Stability** | **Confidence score** | **Stability** |
| **V367F** | -1.121 | Decrease | -0.195 | Decrease | -0.651 | Decrease |
| **R408I** | 0.497 | Increase | 0.007 | Increase | -0.543 | Increase |
| **G476S** | -0.715 | Decrease | -0.45 | Decrease | -0.566 | Decrease |
| **V483A** | -1.175 | Decrease | -0.505 | Decrease | -0.999 | Decrease |
| **N501Y** | -1.067 | Decrease | 0.391 | Increase | 0.790 | Increase |

### Table S5. Protein stability prediction through I-Mutant server

| Position | WT | New | Stability | DDG (Kcal/mol) |
|---|---|---|---|---|
| **367** | V | F | Decrease | -3.08 |
| **408** | R | I | Decrease | -0.21 |
| **476** | G | S | Decrease | -1.52 |
| **483** | V | A | Decrease | -1.03 |
| **501** | N | Y | Increase | 0.15 |

### Table S6. List of residues involved in hydrophobic and hydrophilic interactions during p-p docking.

| | SpikeS1-RBD | | ACE2 | |
|---|---|---|---|---|
| **Variants** | **Hydrophilic** | **Hydrophobic** | **Hydrophilic** | **Hydrophobic** |
| **WT** | Lys417, Tyr449, Ala475, Asn487, Tyr489, Gln493, Gly496, Thr500, Gly502, Val503 | Tyr453, Leu455, Phe456, Tyr473, Gly476, Phe486, Phe490, Gln498, Asn501, Tyr505, | Gln24, Thr27, Asp30, Lys31, Asp38, Tyr41, Tyr83, Thr324, Lys353, Gly354, Asp355, | Glu23, Phe28, His34, Glu35, Leu45, Met82, Asn330 |
| **V367F** | Lys417, Tyr449, Tyr453, Ala475, Tyr489, Thr500, Tyr505, | Leu455, Phe456, Tyr473, Phe486, Asn487, Gln493, Ser494, Gln498, Asn501, Gly502, | Gln24, Thr27, Asp30, His34, Asp38, Thr324, Lys353, Asp355, Ala386, | Lys31, Tyr41, Tyr83, Gly326, |
| **R408I** | Arg403, Lys417, Tyr449, Ala475, Gly476, Tyr489, Gln493, Gly496, Thr500, Gly502, | Tyr453, Leu455, Phe456, Tyr473, Phe486, Asn487, Ser494, Gln498, Asn501, Tyr505, | Glu23, Gln24, Thr27, Asp30, Lys31, His34, Glu37, Asp38, Tyr41, Gln42, Lys353, | Glu35, Leu45, Met82, Asn330, Gly354, Asp355, Arg357, |

| | | | | |
|---|---|---|---|---|
| **G476S** | Lys417, Tyr449, Tyr453, Ala475, Tyr489, Gln493, Tyr495, Thr500 | Leu455, Phe456, Tyr473, Ser476, Phe486, Asn487, Gly496, Gln498, Gly502, Tyr505, | Gln24, Thr27, Asp30, Lys31, His34, Glu35, Asp38, Tyr41, Lys353, Asp355 | Phe28, Leu45, Leu79, Met82, Tyr83, Asn330, Thr324, Gly354, Arg357 |
| **V483A** | Lys417, Tyr449, Asn487, Tyr489, Gln493, Tyr495, Gly496, Thr500, Gly502 | Tyr453, Leu455, Phe456, Tyr473, Ala475, Phe486, Gln498, Asn501, Val503, Tyr505, | Thr27, Asp30, Lys31, Glu35, Asp38, Tyr41, Tyr83, Lys353, | Gln24, His34, Glu37, Gln42, Leu45, Met82, Thr324, Gly354, Asp355, Arg357 |
| **N501Y** | Lys417, Ala475, Asn487, Tyr489, Gln493, Gln498, Tyr501, | Tyr453, Leu455, Phe456, Tyr473, Gly476, Phe486, Ser494, Thr500, Tyr505, | Gln24, Thr27, Asp30, Lys31, Glu35, Asp38, Gln42, Tyr83, Lys353, | His34, Tyr41, Leu45, Met82, Asn330 |

**Table S7. mCSM-PPI Prediction**

| Variants | Predicted Affinity Changes (ΔΔG =Kcal/mol) |
|---|---|
| **V367F** | 0.538 |
| **R408I** | -0.573 |
| **G476S** | -0.058 |
| **V483A** | -0.184 |
| **N501Y** | -2.147 |
